# On prefrontal working memory and hippocampal episodic memory: Unifying memories stored in weights and activity slots

**DOI:** 10.1101/2023.11.05.565662

**Authors:** James C.R. Whittington, William Dorrell, Timothy E.J. Behrens, Surya Ganguli, Mohamady El-Gaby

## Abstract

Remembering events in the past is crucial to intelligent behaviour. Flexible memory retrieval, beyond simple recall, requires a cognitive map, or model of how sensations, actions, and latent environmental or task states are all related to one another. Two key brain systems are implicated in this process: the hippocampal episodic memory (EM) system and the prefrontal working memory (WM) system. While an understanding of the hippocampal system, from computation to algorithm and representation, is emerging, less is understood about how the prefrontal WM system can give rise to flexible computations beyond simple memory retrieval, and even less is understood about how the two systems relate to each other. Here we develop a mathematical theory relating the algorithms and representations of EM and WM by unveiling a duality between storing memories in synapses versus neural activity. In doing so, we develop a formal theory of the algorithms and representations of prefrontal WM in terms of structured, and controllable, neural subspaces (termed activity slots) that together can represent a dynamic cognitive map without any need for synaptic plasticity. By building models using this formalism, we elucidate the differences, similarities, and trade-offs between the hippocampal and prefrontal algorithms. Lastly, we show that several prefrontal representations in tasks ranging from list learning to cue dependent recall are unified as controllable activity slots. Our results unify frontal and temporal representations of memory, and offer a new basis for understanding dynamic prefrontal representations of WM.

## Introduction

Being able to predict what will happen next in novel structured environments constitutes a fundamental feature of intelligent cognition. However, due to the one dimensional and irreversible nature of time itself, we can only learn about structured environments, and the consequences of our actions in them, through a spatiotemporally local sequence of transient sensorimotor experiences. Thus, to learn to predict the future, both brains^1–3^ and machines^4–6^ must convert this transient sequence of sensorimotor experience into internal models of the world that remember and exploit structured relationships between actions, sensations, and environmental features. When such an internal model, or cognitive map, emerges it can allow additional flexibility beyond next step prediction, such as inferring new routes to goals^7^ or simulating counterfactual scenarios^8^. An algorithmic understanding of how local sequential sensorimotor experience is converted into a rich cognitive map capable of predicting the future counterfactual consequences of diverse potential actions remains a major aim of cognitive neuroscience.

Two key brain regions are proposed to build cognitive maps from sequential experience: the episodic memory (EM) system in the medial temporal lobe and the working memory (WM) system in the prefrontal cortex (PFC)^9–11^. However it is not clear why two systems are employed to solve the same problem nor how these brain systems are related in either underlying algorithms or representations.

For sequential episodic memory in the hippocampal formation, ideas are beginning to emerge on underlying algorithms and representations^4,12–14^. In these models, memories are stored in hippocampus (HPC) by updating synaptic connections via Hebbian learning^15^ (like a Hopfield network^16^), with cortical recurrent neural networks (RNNs) controlling which memory to store or retrieve by tracking position (ordinal, spatial, or otherwise) within the sequence. These models are able to explain many cellular recordings in the hippocampal formation for both spatial and non-spatial tasks; hippocampal cells such as place cells^17^, landmark cells^18^, and splitter cells^19^ are explained as memory representations, while entorhinal cells such as grid cells^20^, object-vector cells^21^, border-vector cells^22^, non-spatial grid cells^23,24^, non-spatial sound frequency cells^25^ are explained as representing position, so that the right memory is retrieved at the right position. Importantly, the hippocampal literature shows us that sequence memory is not just remembering sequences exactly as they were presented, but rather using structured knowledge (e.g., where you are in space) to recall the right memory at the right time. Thus even in this case, successful sequence memory goes beyond mere rote memorisation of sensory experience, but rather reflects a more sophisticated cognitive map building process that learns systematic relationships between sensations and actions induced by latent spatial or task structure^4,5,12–14^.

For sequence working memory in PFC, our understanding is limited to tasks of repeating sequences exactly as they were presented. Here, findings from both artificial^26–28^ and biological^29,30^ networks suggest memories of items are organised into neural subspaces according to their ordinal position. Crucially, unlike HPC, storing these memories requires no additional synaptic plasticity since recurrent connections in PFC update and maintain memories stored in neural activity. While this has revealed mechanisms of sequential memory retrieval, PFC is implicated in tasks requiring flexible control of memory retrieval^31–33^, i.e., not just repeating sequences exactly as they were presented. While we don’t understand how PFC working memory models allow such flexible control, we do for HPC episodic memory models. Thus, if we could relate PFC working memory and HPC episodic memory then we could understand how the PFC working memory system flexibly controls its memories to tackle sequence memory and prediction tasks that go beyond simple repetitive sequences and instead require cognitive map-like knowledge of structured relationships between, sensations, actions, and latent spatial or non-spatial variables. Unfortunately no relationships currently exist between PFC working memory and HPC episodic memory. Excitingly, though, recent results from the machine learning literature suggest there may be a relationship. In particular, simplified (linear) versions of transformer neural networks^6^, which are closely related to Hopfield networks^34–36^, have a recurrent reformulation^37,38^ which is suggestive of a relationship between episodic and working memory models.

In this work we provide an understanding of sequence working memory in PFC and how it relates, in representation and algorithm, to HPC episodic memory. In particular we 1) develop a unifying theory of storing sequence memories in synapses (EM) and neural activity (WM); 2) demonstrate that the two algorithms differ in their ability to scale to larger task sizes; 3) demonstrate how the different algorithms utilise different neural representations, with EM using abstract positions, while WM uses slots (distinct neural subspaces) in neural activity; 4) demonstrate that the WM slot representation affords scene algebra; 5) recover single neuron slot coding under certain constraints; 6) demonstrate how internally computed velocity signals can control the contents of PFC slots; 7) show that our theory of controllable activity slots provides a common explanation for PFC data from several disparate studies. Overall, a main dividend of our derived duality between episodic HPC memory and PFC working memory is a new, unifying theoretical framework for how PFC working memory networks could, in principle, implement and control dynamic cognitive maps of the environment or task through recurrent updates of neural activity alone, without any synaptic plasticity.

### Theory: Unifying memories stored in synapses and neural activity

#### Sequence memory and prediction is a structure learning problem

Sequence memory and prediction tasks involve recalling previous observations, but where correct recall depends on an underlying task structure. For example, in Immediate Serial Recall^39^ (ISR), a series of observations are presented in order, and then they must be recalled in order. Here the underlying structure is a ordinal line, but other sequences can be drawn from other underlying structures. For example if a sequence is drawn from navigating in 2D space, then including this structural knowledge in an internal representation will dramatically facilitate recall and prediction, as some transitions are possible in 2D space, but others are not. For example, in a 2D navigation task, it is possible to predict what one will see next when traversing a closed loop for the first time, but this requires not only remembering one’s past sequence of observations, but also combining this sequence memory with knowledge of the structure of 2D space, and how actions move us within the space (i.e. if the integral of your recent velocity will become 0 then you will return to a previously experienced location). This facilitation of sequence memory and prediction by exploiting knowledge of task structure is possible in all problems where there is a common structural constraint. Even when each problem consists of sequences with different observations (so one problem’s sequence cannot be memorised and used for another problem), knowledge of the common task structure across problems facilitates recall and prediction within each problem. In machine learning terminology, the underlying task structure must be meta-learned across problems (learning to learn; Figure 1A left for example ISR task).

**Figure 1.**
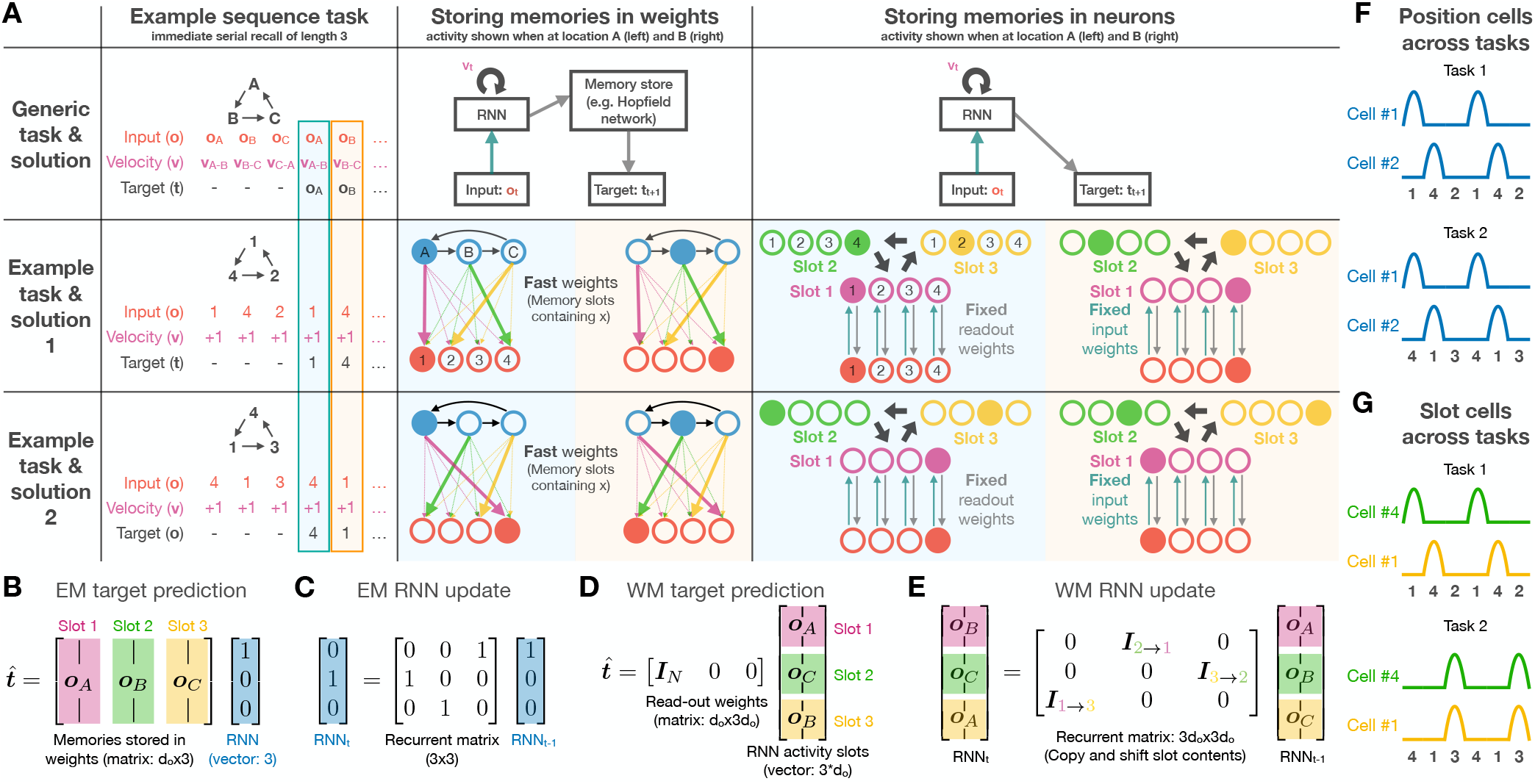
Relationship between episodic memory and working memory models. **A)** Schematic of how a task can be solved by episodic and working memory solutions. Solid/empty circles are active/inactive neurons. Black/teal/grey arrows are recurrent/input/output connections. Left/Center/Right: Task/EM solution/ WM solution. Top/Middle/Bottom: General example/ Specific example 1/ Specific example 2. **Left:** Example task of Immediate Serial Recall (ISR) of length 3, with specific instances below. The task input is a sequence of 3 random observations, ***o***, that repeat, and a constant ‘forward’ velocity, ***v***. The task is to predict the upcoming target, ***t***. Blue/orange segments highlight the timesteps described in EM/WM solution schematic (center/right). For the purposes of the schematic, we assume each observation has a one-hot encoding. **Center:** Top: EM models involve an RNN that indexes memories stored in synaptic weights. Middle/Bottom: Schematic of EM solution. This is a simplified version of the top figure (e.g., the RNN is one-hot, and memories are stored in synapses between the RNN and observation neurons; full model and derivation of simplification in Appendix A.1), but it maintains all the key elements. Past observations are stored in synaptic memory slots (here we show synaptic memory slots as non-overlapping sets of neural weights), and recurrent position representations index memory slots to retrieve a memory for predicting the upcoming target. Red neurons one-hot encode observations (each neuron represents one observation: 1, 2, 3, or 4). Blue neurons one-hot encode position (each neuron represents one position: A, B, or C). Blue/orange segment highlights present the network at two different timesteps of the task. **Right:** Top: WM models involve just an RNN. Middle/Bottom: Schematic of WM solution. Past observations are stored in activity slots within the RNN (we show activity slots as non-overlapping sets of neurons, but they could be non-overlapping neural subspaces more generally), with the content of each slot copied and shifted by the RNN (at every timestep) to successfully predict targets using fixed readout weights. This example RNN has three activity slots (Green/yellow/purple) each consisting of 4 neurons that can one-hot encode 4 possible observations (1,2,3,4). Blue/orange segment highlights present the network at two different timesteps of the task. **B-E)** Mathematics of the EM and WM solutions. **B)** The EM solution makes predictions via an RNN position code (blue vector) that indexes memories stored in synaptic weight slots (matrix columns). **C)** The RNN position code is updated by recurrent connections. **D)** The WM solution makes predictions using fixed readout weights that attend to a single RNN activity slot (slot 1). **E)** Contents of RNN activity slots are copied and shifted by a recurrent matrix. Note the recurrent matrices will be different depending on the task structure. **F)** The activity of two example position neurons (cells #1 and #2 are the left/middle blue neurons in panel A middle). Cell #1 / #2 is active at position A/B. Thus the two neurons have similar activity patterns in both tasks (top/bottom), up to a possible global shift in their firing fields across tasks, as absolute position could be unknown. However, the neurons preserve their *relative* phase relationship across tasks; i.e. the phase lag between the activity peaks of cell #1 and #2 is preserved across all tasks. **G)** The activity of two example activity slot neurons (cell #4 is rightmost green neuron, and cell #1 is leftmost yellow neuron in panel A right). Cell #4 always fires 2 steps after observation 4. Cell #1 always fires one step after observation 1. In task 1, these neurons have the same activity as the above position neurons, but do not in task 2: they do not keep their phase relationships between tasks as they each code for specific distance/lag from specific observations.

#### Task and problem formalism

Formally, for each task, we consider a dataset *𝒟* = {*𝒟*_0_, *𝒟*_1_,*…, 𝒟*_*N*_ }. Each *𝒟*_*i*_ is one of *N* problem instances of the common task, and is simply a sequence consisting of vectors of sensory observations, ***o***, of dimension *d*_*o*_ (termed 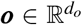), sensory targets, 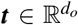, and (allocentric) velocities, 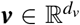, i.e., *𝒟*_*i*_ = (***o***_0_, ***t***_0_, ***v***_0_), (***o***_1_, ***t***_1_, ***v***_1_),…, (***o***_*K*_, ***t***_*K*_, ***v***_*K*_) if the sequence is of length *K*. These velocities reflect actions taken by agents, but in this case we provide them externally. Importantly, the underlying structure of the task is determined by how successive actions cumulatively change latent states within the task, which themselves are never *directly* observed by the agent.

Concretely, we assume there is an underlying latent variable, 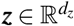, which we can think of as corresponding to a latent agent or task state. In spatial contexts we can think of the agent’s latent state as its position. Furthermore, we assume each latent state, or position ***z***, is associated with a visible observation ***o***, corresponding to what the agent would sense if it had the latent state, or position ***z***. Also, we assume that for any ordered pair of neighbouring source and target latent states, there is a unique action, or velocity ***v*** that modifies the agent’s latent state from the source to the target (see Figure 1A left for a simple loop structure with one action). The task structure is encoded by how actions, or velocities, change the latent state, or position. For example, in the case of navigation in a 2D grid, there are 4 elementary actions, or velocities, corresponding to moving one step `North`, `South`, `West` or `East`, and each step moves the latent 2D grid position ***z*** by one step in the corresponding direction. In our formulation, successive actions correspond to the addition of velocities, and so when a sequence of velocities additively cancel (i.e. `North` + `East` + `South` + `West` = 0 for a 2D grid), the then the agent necessarily returns to the same latent position and encounters the same observation at the beginning and end of the action sequence.

Now, returning to the data for a single task *𝒟* = {*𝒟*_0_, *𝒟*_1_,*…, 𝒟*_*N*_, }each individual problem instance *𝒟*_*i*_ assumes a different and random pairing of observations ***o***, to latent states ***z***. However, all *N* problem instances share the *same* underlying task structure, specified by how actions, or velocities ***v*** change latent states. The aim of the task is to predict a target, ***t***, at each timestep which is either an upcoming observation (i.e., what you will see after going `North`), or a past observation (i.e., what you saw 5 steps ago). Importantly, neither latent states, nor how actions affect latent states, are directly observed by the agent. Instead the agent only experiences a spatiotemporally local sequence of transient sensations and actions. To successfully solve the prediction or recall task across all problems, the agent must build a cognitive map of the common task structure that captures how actions affect latent states, and how latent states are bound to observations. Only then, for example can the agent predict what it will see next upon the first traversal of a closed loop in a 2D grid. Finally, we note that while the task formalism (and subsequent model formalism) is general to tasks with discrete and continuous positions, we primarily consider the discrete setting here (with *n*_*p*_ total positions).

Next we will develop a theory for a very simple network realisation of an EM solution, and from that derive a very simple network realisation of a WM solution. But crucially, we will show below that many of the features of these simple solutions hold in more general scenarios, both in generic artificial networks of machines, and in biological networks of animals, that successfully solve structured sequence memory and prediction tasks.

#### Solving sequence memory tasks with EM

The hippocampal literature has developed EM models that solve these tasks^4,14^. These models are built from two components (1A top-middle). The first component is an RNN that learns to *explicitly* represent position in a manner that generalises across tasks. The second component is a memory system that binds observation representations to position representations and stores this memory in synaptic weights. Critically these weights change in every problem so different problems from a common task can be solved with a common position code. In its simplest form, the RNN has a single neuron active for each position (Figure 1A middle; full theory in Appendix A.1). Sensory memories for each position are encoded in synaptic weights between each RNN neuron (corresponding to that position) and the observation representation, with memories added (weights updated) via Hebbian learning. The role of the RNN is to track the underlying latent variable, ***z***, so that the correct position cell is activated at the right time to retrieve the right memory (Figure 1A middle; the leftmost position cell activates to recall the observation at position A). Tracking position means the RNN integrates velocity signals, ***v***, to update its representation from ***h***(***z*** − ***v***) to ***h***(***z***) (i.e. from position ***z*** − ***v*** to position ***z***). This can be done with velocity dependent matrices, ***W*** _***v***_, that follow the transition rules of ***z*** (e.g., ***W***_***v***_***W*** _−***v***_ = ***I***; going `North` then `South` takes you to the same position). Mathematically, the RNN update is ***h***(***z***) = ***W*** _***v***_ ***h***(***z***− ***v***) (Figure 1C). The overall process of generating target predictions (assuming all observations have already been seen and added as memories - the ***o*** terms in the below equation) is succinctly described in the following equation:

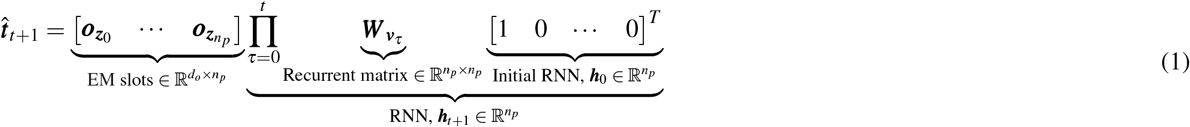

We see that, an initial RNN representation, ***h***_0_, is successively updated by velocity dependent matrices, ***W*** _***v***_, to code for position at timestep *t* + 1: ***h***_*t*+1_. To make predictions, ***h***_*t*+1_ selects (like an attention vector) one of the *n*_*p*_ observations stored in memory **slots** in synaptic weights (Figure 1B; synaptic memory slots are non-overlapping sets of synaptic connections that can be rapidly updated to store arbitrary memories; each slot corresponds to a position ***z*** and stores the observation at position ***z***; ***o***_***z***_). For clarity of presentation, we chose a basis in which the RNN vector is one-hot (has one cell active at any time) and the matrices ***W*** _***v***_ have columns that are all zeros except for a single 1 (e.g., Figure 1C), though the above solutions works in any basis, i.e., ***h***← ***O***^*T*^ ***h, W*** _***v***_ ← ***O***^*T*^ ***W*** _***v***_ ***O***, etc. Indeed, under simple constraints, the optimal solution to this basis is a grid cell basis^4,40^.

#### Solving sequence memory tasks with WM

The above solution stores memories in synaptic connections, and is thus relatable to the hippocampal formation. However, the PFC is thought to store sequence memories in the dynamics of neural activity as opposed to synaptic connections^27,30^ (1A top-right; although see^41^). Here we show that reshaping and rearranging the above equation produces an alternative, but equivalent, solution where memories are stored in RNN activity rather than synaptic connections (Figure 1A right):

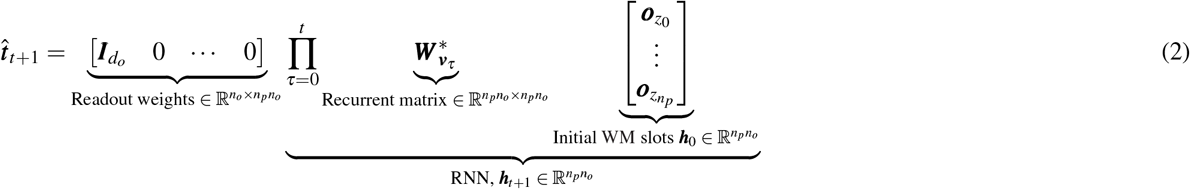

This equation performs the exact same computation as the previous equation, though its terms, while similar, have important differences (see Table 1 for relationships between terms). In particular, what was a matrix of memories stored in synaptic weights, 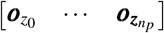, is now a vector of memories stored in RNN activity, 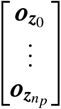 Thus the total neural activity pattern can be decomposed into a collection of non-overlapping subspaces which we term **activity slots**, and each such slot can store a memory of an arbitrary observation (Figure 1A right). Readout weights are now fixed and attend to a single slot (Figure 1D). Thus, to readout a memory, the contents of each slot must be copied and shifted to other slots via recurrent weights 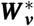, so that the correct observation can be readout (in the above equation the first slot is the readout slot, but in general it could be any slot - this is set by the structure of the task, e.g., Figure 8). To copy and shift the contents of activity slots, the WM velocity dependent matrices, 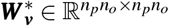, are expanded (and transposed) versions of the EM velocity-dependent matrices, ***W*** _***v***_: where there was a 1 or a 0, there is now an identity matrix 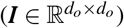 or a matrix of zeros 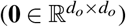 (See Figure 1E for a schematic of the structure of ***W*** _***v***_). Intuitively, the matrix is bigger (by a factor of *n*_*o*_ in both dimensions) as we are now simultaneously tracking relative position to each observation, rather than abstract task position. Adding a memory to a slot is simple. Feed-forward weights can direct observations directly to the input slot (e.g., Figure 1A right). The memory is then passed from slot to slot, controlled by velocity. When memories reach the output slot, it can influence immediate behaviour. Thus, once learned, this solution requires no synaptic plasticity. Lastly, similar to the EM solution, the WM solution works in any basis (not just the one presented above).

**Table 1.**
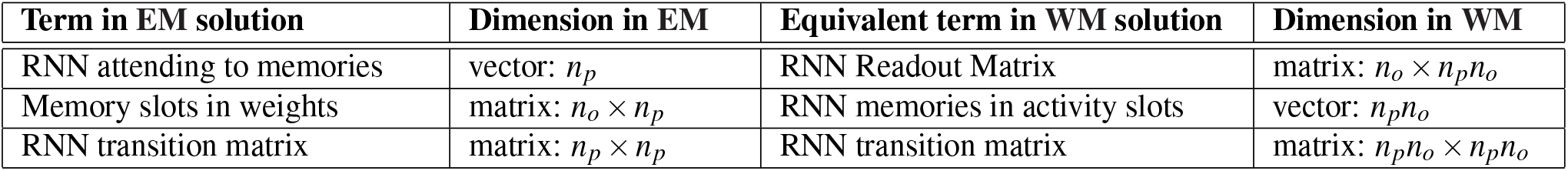
Equivalent terms in the EM and WM formulations, along with their dimensions. We see that vectors (activity) turn into matrices (synaptic weights), and vice versa. *n*_*p*_ is the number of positions in the task, *n*_*o*_ is the dimension of the observation vector (***o***).

#### Memory slots unify episodic and working memory and yield a rich neural basis for WM cognitive maps

While both solutions use memory slots, they do so in conceptually and functionally different ways. In the WM solution, memory slots are structured neural subspaces, with memories for different observations stored in each neural subspace, and the contents of each subspace can be copied and shifted to other subspaces. In the EM solution, memory slots are in synaptic connections (each memory is stored in a subspace of the weights), with each memory statically remaining in its slot (subspace). The memory slots are also accessed differently. The EM solution uses flexible attention (via an RNN) to index ‘fixed’ memory slots (the weights don’t move), while the WM solution uses fixed attention (the readout weights) but has flexible memory slots (the contents can be copied and shifted by the RNN)^i^.

Interestingly, there is a striking representational difference between the WM and EM solutions. This difference arises because the RNN in the EM solution represents latent position alone (relative to some initial position), independent of sensory observations attached to any position in any problem or task. However, RNN neurons in the WM solution represent relative position *from* individual observations - a type of conjunctive representation of relative position *and* observations for every problem. In particular, the slot identity corresponds to a particular relative position from the agent’s current position (or more generally a particular sequence of action/velocities one could take in a structured environment), and the contents of that slot is the observation at that relative position. In other words, the contents of each slot indicates the correct counterfactual answer to the task *if* the slot’s corresponding action *were* taken. For example, in the simple case of Figure 1A, third column, Slot 1, aligned with the input/readout weights, contains the correct answer to the prediction task if no action were taken. And Slot 2 and 3 contain the correct counterfactual predictions if the agent *were* to go back one or two positions respectively. Every time the agent takes an actual action, the correct counterfactual prediction associated with each slot necessarily changes, and so the WM RNN must update the contents of very slot by appropriately copying and shifting their contents through the recurrent weights. Because of this dynamic update, at any given time, the entire WM representation acts like a cognitive map that can *simultaneously* answer 3 counterfactual questions (what observation would I predict if went back 0, 1, or 2 positions) using decoders that attend to slots 1, 2, and 3, respectively. As the agent moves forward in position, the slot activity is shifted backwards correspondingly so that *each* slot always contains the correct answer its *own* action specific counterfactual question. Moreover, in the simple case of Figure 1A, third column, since each neuron in each slot is selective for a single observation, from a single neuron perspective, the neural code will consist of a collection of neurons that fire selectively for particular observations, at particular times since the observations were most recently encountered (or more generally for particular observations and particular relative positions / action sequences taken since the observation). Any single neuron’s relative position / action selectivity elegantly derives from the *identity* of the slot it is a part of, while that same neuron’s object selectivity derives from its functional role *within* its slot.

This fundamental difference between EM and WM solutions implies that single neurons behave very differently across tasks. In particular, any two EM position neurons that fire next to each other in one task, will fire next to each other in another task: they maintain their phase relationship (Figure 1F; just like grid cells do). On the other hand, two activity slot neurons with a particular phase relationship in one task (Figure 1G, top), may not have the same phase relationship in another task (Figure 1G, bottom). This is because WM cells code for relative position (as defined by the slot’s intrinsic relative position to the input/readout slot) to a particular observation (as defined by the identity of the neuron within a slot). Since the pairing between positions and observations are shuffled between different problem realisations of a given common task, the relative phase between WM cells will also appear shuffled. Interestingly, as we will see below, this slot derived conjunctive coding of observations and actions, reflecting a cognitive map that can simultaneously answer many counterfactual questions contingent on different actions, can elegantly and succinctly account for a wide range of otherwise perplexing properties of rich prefrontal cortical representational dynamics, both *in silico* and *in vivo*.

### Model architectures and task specifics

To experimentally test the predictions of the theory, namely that the EM system uses position representations whereas WM systems use activity slots, we use a variety of tasks and model architectures (full details in Appendix A.2).

#### Model architectures

We build both EM and WM neural network models (Figure 2A,B). The key difference between the two models is that the EM model makes predictions by retrieving memories from an external memory system, whereas the WM model makes predictions with a learned readout matrix. For the EM external memory, we use a modern Hopfield network^35^, which our above theoretical results are derived from (Appendix A.1). Both models use an RNN to control memories. To be general, we use two RNN variants (Figure 2C,D); one directly inspired by our theory that uses velocity dependent transition matrices (GroupRNN; related to selective state-space models^42^), and the other is a conventional RNN which receives velocity signals as input (RegularRNN). All model parameters are initialised randomly, and trained using backpropagation.

**Figure 2.**
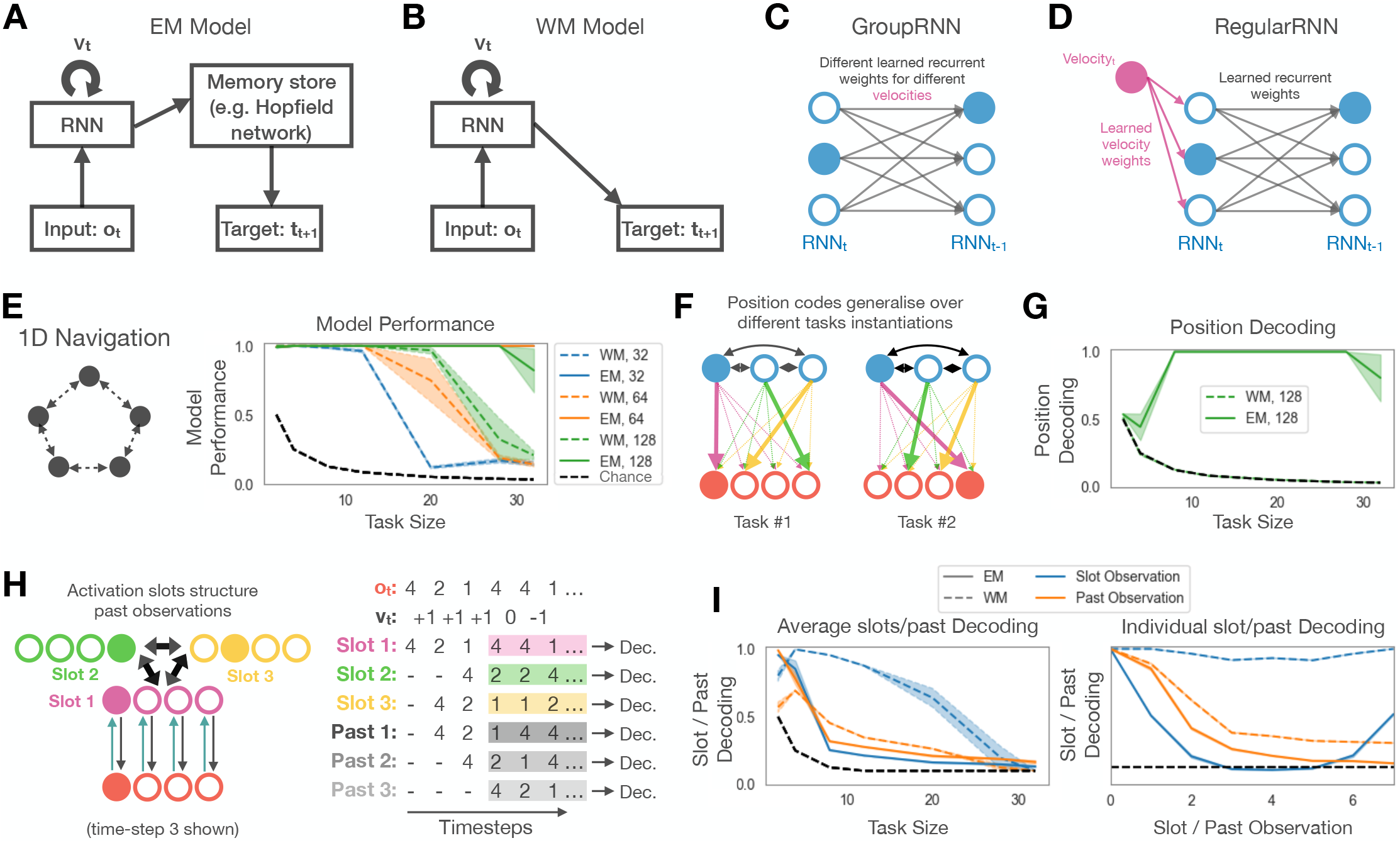
EM models scale better and learn position representations, while WM models learn activity slots. **A)** We train EM models consisting of an RNN that can store and retrieve memories. At each timestep, the RNN receives an observation, ***o***_*t*_, and velocity signal ***v***_*t*_, and must predict target ***t***_*t*+1_. **B)** We train WM models that are just an RNN. We use two types of RNN (for both EM and WM models): **C)** Like our theory we use an RNN with learnable velocity dependent matrices (GroupRNN); **D)** We also use a conventional RNN which receives velocity as input (RegularRNN). **E)** Performance of the EM (solid) and WM (dashed) models on 1D navigation tasks (using GroupRNN). Black dashed line denote chance performance. EM models perform better that WM as task size increases. Larger sized RNN perform better for both EM and WM models. For each combination of task, task size, model type, and RNN size (32, 64, 128), we train five randomly initialised models. Results are mean *±* standard error. All subsequent panels are for RNNs of size 128. **F)** Our theory says EM models should learn position representations that generalise over task instantiations (i.e., in sequences with different sensory observations but same underlying structure). **G)** EM models, but not WM models, learn position like representations. **H)** Left: The WM activity slot theory says the RNN organises past observations into slots (structured according to underlying task structure). Right: Thus a particular sequence of observations should be decodeable for each slot (we call this the ‘slot-sequence’ for each slot). This sequence is different to the sequence of time-lagged past observations (we call this the ‘past-sequence’ for each time lag). **I)** Left: Average decoding of ‘slot-sequences’ (what observation should be in each slot according to our theory) and ‘past-sequences’ (what observation occurred *i* timesteps ago, up to the total number of slot). WM models have high slot-sequence decoding, but not past-sequence decoding, indicating they have learned activity slots. The EM models have not learned activity slots, apart from those trained on small task sizes. Right: Slot- and past-sequence decoding for individual slots or time lags for 1D navigation on an 8-loop. Slot-sequence decoding is consistently higher than past-sequence decoding for WM models, but not EM models, for all slots/time-lags. Similar results shown for all tasks with GroupRNN in Figure 11, and for all tasks with RegularRNN in Figure 12.

#### Tasks

We consider four main tasks (Figure 8A) that are all related to the neuroscience and cognitive science literature (further neuroscience tasks are introduced later): Immediate Serial Recall (ISR), N-Back, 1D Navigation, and 2D Navigation. These tasks range in complexity from simple sequence repetition that requires no velocity integration to tasks where correct recall relies upon integration of progressively more complex velocity signals. In all tasks, observations, ***o***, and velocity signals, ***v*** are provided at each timestep and the models are trained to predict a target, ***t***, at each timestep. After observing the observation and velocity at timestep *t*, the model is asked to predict the target at timestep *t* + 1. An example problem sequence for the 1D Navigation task (a random walk on a loop; e.g., a 3-loop with observations 1, 3, and 2 at the three positions) is: ***o*** = {1, 3, 2, 2, 3, 1, 2, … }, ***v*** = {+1, +1, 0, − 1, −1, − 1, … } and ***t*** ={ −, −, −, 2, 3, 1, 2, … }. Here, ‘−’ means a target that we do not train on since it has not been observed before and therefore is not possible to predict. The ISR task is like 1D navigation but with constant forward velocity, the 2D navigation task is a 2D version of the 1D navigation task, and the N-Back task is a random observation sequence (with constant velocity signal) and the targets are observations N timesteps ago. For each task we train on multiple problem realisations corresponding to multiple sequences where position-observation pairings are randomised, but the underlying task structure of how actions affect positions (e.g., 1D Navigation) is fixed. See Appendix A.3 for details on all tasks. It is important to note that the 1D and 2D navigation tasks (as well as some tasks later in this paper) involve more than just repeating the sequence exactly as it was presented and so cannot be solved without learning a cognitive map and correctly integrating velocity signals.

## Results

### Differences in EM and WM algorithm performance and representation

#### EM scales more favourably than WM

Our theory demonstrates that EM and WM systems implement the same computation (solve the same task) but use different underlying algorithms (their neural mechanism is different). These algorithms have different trade-offs. Most notably, the WM solution requires many more neurons than the EM solution (a factor of *n*_*o*_ more since each observation must be stored in neural activity). Thus we predict that, for a fixed neuron budget, the WM model performance will degrade when the number of memories needed to be stored, which we refer to as the task size, increases, while the EM model will continue to perform well. We demonstrate this phenomenon for the 1D navigation task (Figure 2E) as well as the other tasks (GroupRNN: Figure 11A, RegularRNN: Figure 12A). Additionally, we observe performance improves when the WM RNN has more neurons, as predicted. Also, in both EM and WM models, we observe reduced performance in tasks with more action dimensions (e.g. 2D navigation versus ISR; Figures 11A, 12A), likely because more weights need to be precisely tuned with more action dimensions.

#### EM uses position representations while WM uses activity slots

Our simple theory above predicts that even more potentially complex trained EM models should use position-like representations to index memories stored in synaptic weight slots (equation 1), whereas trained WM models should store and manipulate memories in RNN activity slots (equation 2). To test these predictions, we train linear decoders to decode positions and observations from the neural activity of trained models’ RNNs. Crucially, we avoid task over-fitting by training each decoder on many example tasks, rather than just one, demonstrating that the activity slots, or position representations, are generalisable across tasks.

First, our theory says abstract position (independent of object identity) should be decodeable from EM models, but not WM models, since EM models use position to index memories (Figure 2F). We confirm this prediction in all tasks and for all RNN types (Figures 2G, 11B, 12B). This is consistent with existing EM models^4,14^ that learn position codes to index hippocampal memories, as well as entorhinal data such as grid cells that code for position. We note, however, that position is often not decodable for large or small task sizes. For large sizes, it is easy to understand: the models were unable to learn the task and thus learned no sensible representation (Figure 2E, 11A, 12A). For small task sizes, the RNN in the EM model has learned to use another memory indexing representation, namely the WM solution. We discuss this case further in a later paragraph.

Second, our theory predicts that previous observations should be decodeable from the RNN in WM models, but not the EM models, since the RNN in WM models store memories of previous observations in neural activity (Figure 2H) while RNN activity in EM models only tracks position, with memories of previous observations stored in synaptic connections. More precisely, our theory prescribes exactly which past observation is stored in which neural activity slot at any given time since it details how slot contents are copied and shifted (or more generally rotated) by the trained RNN connectivity in a manner that reflects the underlying structure of the task (i.e. how velocity signals change position) (Figure 8B,C). Thus, over time, the neural activity within each activity slot will encode a particular sequence of observations. Moreover, for tasks with varying velocity signals, this sequence will not be a simple temporally shifted copy of the input observations (Figure 2H). We refer to this sequence, defined purely by the task structure, velocity signals and observations, as the slot-sequence for each predicted activity slot. We find that these specific slot-sequences can indeed be successfully decoded from RNN activity in all tasks for WM models (Figures 2I left, 11C, 12C), whereas other highly natural past-sequences (e.g., the observation *N* steps ago) cannot be decoded (apart from the ISR and N-Back task where past-sequences and slot-sequences are identical; Figures 11C, 12C). Slot-sequence decoding performance degrades for larger task sizes, as the RNN fails to learn the task. Furthermore, for an example task size on each of our four tasks, we show high individual slot-sequence decoding for WM models (Figures 2I right, 11D, 12D). These results demonstrate that WM models learn activity slots, and they use velocity signals to copy and shift previous observations between slots: velocity signals control activity slots.

Interestingly, while the EM models have, as predicted, poor slot-sequence decoding and high abstract position decoding in most situations, for small task sizes the converse can be true (Figure 11G,I). This is because an activity slot representation can also be effective at indexing memories; like a position representation, activity slots uniquely code for each position in each task (however it does not code position the same way across tasks, hence we cannot decode *abstract* position from activity slots). We posit the WM solution is easier to learn and hence preferred when possible (i.e., when not limited by number of neurons), which intuitively makes sense as slot representations are a direct function of input data, while position representations are abstract.

### Visualising slots & slot algebra

#### Visualising slots without decoding

We have so far provided evidence, through decoding analyses, that WM models learn activity slots, as predicted by our theory. We focused on decoding analyses because learned activity slots will in general correspond to different neural activity subspaces, that are not necessarily aligned to single neuron axes; i.e. single neurons could exhibit mixed selectivity for multiple slots. To more directly visualise learned activity slots in an even simpler setting, we leverage a recent regularization technique^43^ that encourages demixed representations in which single neurons code for single independent factors (e.g., in our case slots). In particular, we train models while enforcing RNN activity to be nonnegative (i.e., a ReLU activation function rather than linear or tanh) and energy efficient (i.e., minimum norm weights and activities).

With these constraints, single neurons do indeed participate only in a single slot (Figure 3A). We quantify the degree to which single neurons code for individual slots, using the metric ‘mutual information ratio’ (MIR), a well grounded metric^43,44^ where high values indicate demixed single neuron slot coding. We demonstrate MIR improves with the additional constraints across all four tasks and for both RNNs types (Figure 13; GroupRNN and RegularRNN; blue and orange colours).

**Figure 3.**
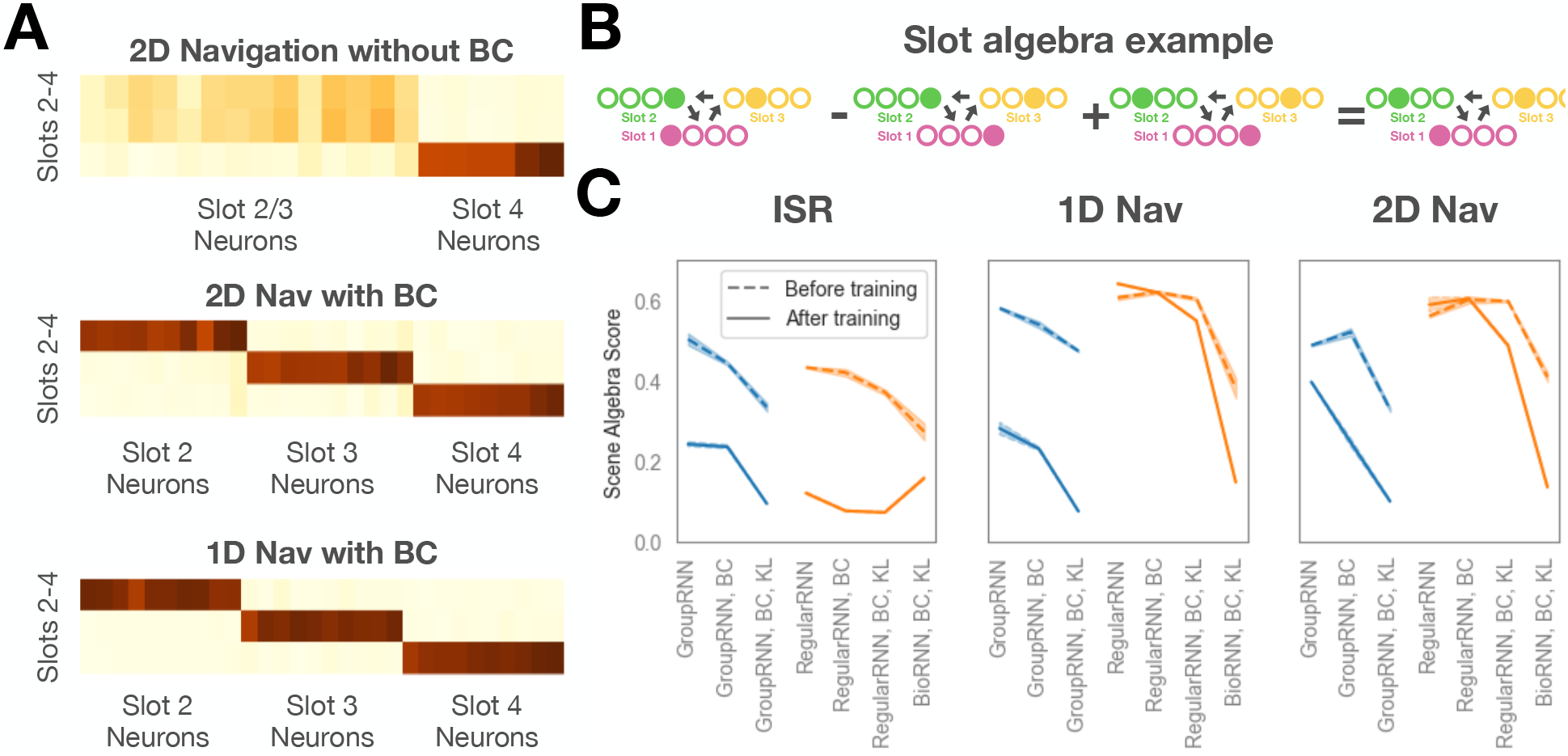
Visualising WM slots without decoding, WM models afford slot algebra. **A)** To visualise WM activity slots we train models while enforcing non-negative RNN activity (using a ReLU) as well as regularising RNN activity (BC: a technique shown to encourage single neurons to code for single factors of variation^43^). We show the mutual information matrices between neurons (horizontal) and slots (vertical). We only show neurons with non-negligible activity, and only present slots 2-4 for clarity since slot 1 tends to have many more neurons than slots 2-4. Top: Without the technique, single neurons code for multiple slots. Middle/Bottom: With the technique single neurons code for single slots. **B)** Slot algebra means representations from three different tasks can be combined to give the representation of a fourth task, e.g., *Rep*_*i*_([1, 4, 2]) − *Rep*_*i*_([4, 4, 3]) + *Rep*_*i*_([4, 2, 3]) = *Rep*_*i*_([1, 2, 2]), where *Rep*_*i*_([*a, b, c*]) denotes the RNN representation at position *i* in a task with observations *a, b, c* at each position. The slot algebra score is a measure of this property, with lower values indicating more accurate implementations of this property. **C)** WM models display ‘slot algebra’ with slot algebra scores low for all GroupRNN models (blue) after training, and are low for RegularRNN models when using the BioRNN variant. For each task and model type, we train three randomly initialised models. Results are mean *±* standard error.

#### Slot algebra predicts relations between neural representations of different problems from a common task

To further demonstrate that RNNs learn activity slots, as well as show that activity slots permit compositional computations, we test another prediction of our theory: activity slot representations should obey a slot algebra, which predicts remarkable and simple linear relationships between WM representations corresponding to *different* problem realisations of the same underlying task. As an example, consider a task with 3 latent positions and 4 possible observations (Figure 3B). Let *Rep*_*i*_([*a, b, c*]) denote the entire WM RNN representation while the agent is at position *i* in a problem realisation with observations *a, b, c* paired at each of the 3 positions respectively^ii^. Now consider the neural representations for two problems, *Rep*_*i*_([1, 4, 2]) and *Rep*_*i*_([1, 2, 2]). These two problems differ only in that the observation 4 at position 2 is replaced with the observation 2. The replacement of one observation with another induces an action on neural representations yielding the slot algebra relation *Rep*_*i*_([1, 2, 2]) = *Rep*_*i*_([1, 4, 2]) − *Rep*_*i*_([*a*, 4, *c*]) + *Rep*_*i*_([*a*, 2, *c*]) (Figure 3B illustrates this relation when the agent is at position *i* = 0). Here *a* and *c* in the middle two terms can be any observations; the key is that the algebraic relation corresponds to taking away observation 4 at position 2 via subtraction of neural representation *Rep*_*i*_([*a*, 4, *c*], for any *a* and *c*, and replacing it with observation 2 via the addition of another representation *Rep*_*i*_([*a*, 2, *c*]) with the *same a* and *c*. Critically, this equality should be true independent of the particular sequence/trajectory taken to position *i*.

To test whether our model representations obey slot algebra, we examine the WM RNN in situations that satisfy the equality relation above. Each relation involves neural representations from 4 separate runs of the RNN in 4 *different* problems. We sum the 3 representations according to the right hand side of the relation, and compare the sum to the measured representation on the left hand side using a squared difference measure (*d*_1−2+3−4_). We then compare this measure to the squared difference of the first term on the left and the first term on the right (*d*_1−2_) via the ratio 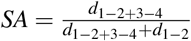. Low *SA* values indicate the representation obeys slot algebra.

The correctness of slot-algebra requires trajectory independent representations, that depend only on the current agent position independent of how the agent arrived at the position. However, not all methods of integrating velocity in neural networks are trajectory independent, with some attractor networks, both in biology^45^ and models^46^, having velocity modulated neurons. Indeed it is possible to have a slot-based representation where slots are conjunctive with velocity. Thus, understanding whether neural representations obey slot-algebra, not only assesses compositionality, but is also informative of how the RNN integrates velocity. Here we test which RNNs obey slot-algebra^iii^. We test the GroupRNN which modulates synapses with velocity signals (and is the model analogue of our theory), the RegularRNN which adds velocity signals to its RNN activities, and, additionally, the BioRNN which integrates velocity signals in a separate neural population, but not the RNN activities themselves. As predicted, the GroupRNN has low *SA* for all datasets (Figure 3C) since velocity does not directly modulate RNN activity. *SA* further improves with additional regularisation (KL) that further encourages representations to be consistent across timesteps, i.e., trajectory independent (see Appendix for details). Conversely, as predicted, the RegularRNN only achieves low *SA* on the ISR dataset which requires no velocity signal (see Figure 10 for another RegularRNN variant which exhibits the same phenomena). Lastly, the BioRNN achieves low *SA* for all datasets. Intriguingly the architecture of the BioRNN is closely related to the neural circuit discovered in the fly head direction system^47–49^ (Figure 7E,F; details in Appendix A.2).

### Activity slots controlled by velocity signals provide a unifying framework for diverse PFC representational dynamics

We have shown that our theory of controllable WM activity slots explains representations that emerge in artificial RNNs trained on structured sequence memory and prediction tasks. The implication of this is that any task where latent positions are linked through action induced transitions can be optimally navigated using WM controllable activity slots. This, however, relies on the appropriate integration of velocity signals. A key question that remains is what are these velocity signals, and how are they represented in neurons? In this section, we demonstrate that PFC-dependent sequential memory tasks and cued memory tasks can be unified through activity slots, but where the distinguishing feature is the velocity signal that controls activity slots.

#### PFC and RNNs compute and represent velocity signals

In previous tasks, we directly provided velocity signals to the model (thus specifying its representation). However, actual brains do not have this luxury; they must compute velocity signals from lower level sensory inputs and choose how to represent them. In spatial tasks, self-movement vectors are thought to be computed from vestibular or other sensory inputs and represented using head direction cells^50^ and speed cells^45^. For complex PFC tasks, where behaviour may be non-spatial, it is unknown what the velocity signals are, or how they are represented.

Recent neural data however, while not demonstrating velocity signals, have found neurons that track progress to goals, invariant to the actual distance to the goal^30,51^. For example a 50% ‘progress cell’ fires at the midpoint between two goals whether they are 3 steps apart or 20. In order to update progress, we contend ‘progress-velocity’ signals must be computed and represented. To test for this, we train WM models on analogous tasks, but to be general and illustrate the principles of working memory, we remove the spatial component to these tasks.

In particular, Basu et al.^51^ trained rodents to run back and forth between two (of 9) reward ports, and El-Gaby et al.^30^ trained rodents to repeatedly visit 4 (of 9) spatially located reward ports in sequence. In both studies, the animals are trained on many problem realisations with random goal positions. We model these tasks (modulo the spatial component), as a 2-ISR and a 4-ISR task, but with delays between each observation (Figure 4A,E left; just like the ‘delay’ of travelling between two spatial goals). Critically, while the delays are random across different problem realisations of each task, the delays are the *same* on each loop on every problem. In a spatial context, this is analogous to how distances between goals are the same on a particular task, but change for different goal positions on new problem realisations of the task. For example, possible 2-ISR delay sequences for different problem realisations are [4, *, *, 2, *, *, 4, *, *, 2, …] or [1, *, 3, *, 1, *, 3, …] where is the common delay observation (that also must be predicted), and possible 4-ISR delay sequences are [4, *, 2, *, *, 2, *, 6, 4, *, 2, *, *, 2, *, 6, 4, …] or [1, 3, *, 8, *, 2, 1, 3, *, 8, *, 2, 1, …]. We provide no explicit velocity signal, and so the model will have to utilise the sparse inputs to compute an appropriate progress-velocity for each delay period, and remember it on returning to that same delay (i.e., after0020one full loop of experience).

**Figure 4.**
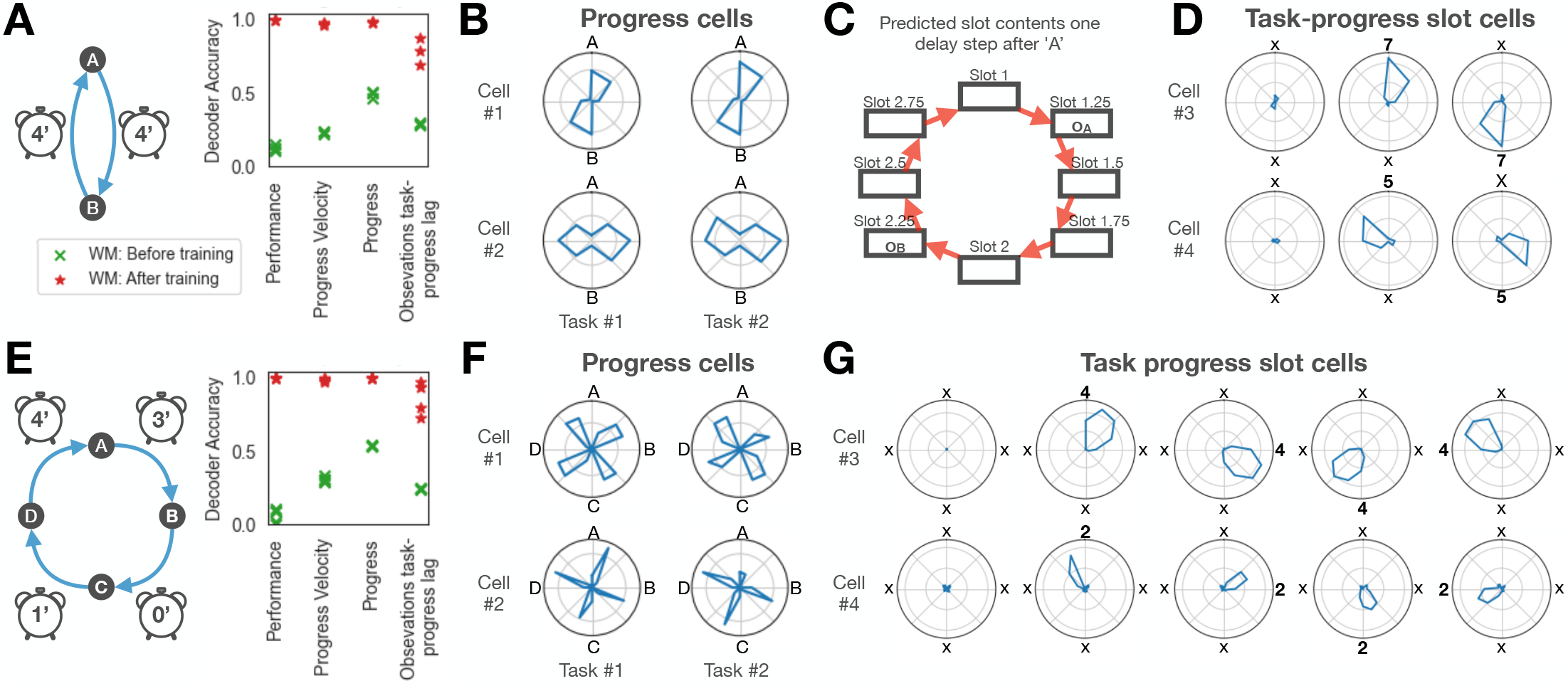
WM-RNNs computes velocity to control activity slots. **A)** We train an RNN on a 2-ISR task, but with delays between observations. Delays are sampled randomly from {3, 4, 5, 6, 7} for each problem realisation of each task, and are fixed for the duration of the individual problem. An example delay of 5 steps shown in the schematic. An RNN is trained on multiple problems simultaneously, and so, after training, must be able to learn about and exploit the random delay in each new problem directly through experience, without any plasticity. Decoding analyses reveal that progress-velocity (reciprocal of delay duration) and progress (% of delay length completed) are represented, i.e., the models maps varying delay lengths across problems into a common progress representation. We can also decode observations at the predicted ‘task-progress’ lag (e.g., ‘3’ was 125% progress lengths ago) according to our model of task-progress activity slots. Red/green points correspond to before/after training. **B)** The RNN learn progress cells, i.e., cells that fire at a consistent progress post observation (regardless of observation type). Angle of circle represents fraction of progress from goal A at 0 degrees top to goal B at 180 degrees bottom, and then back. Radius of blue curve indicates firing rate of cell as a function of fractional progress. The bimodal firing rate indicates the cell fires at a fixed fraction of progress to the next goal, independent of goal identity A or B. These cells transform the variable delay periods for each problem into a common representation. **C)** Schematic of task-progress activity slots. Rather than two slots for this 2-ISR task with delays, there are multiple intermediate slots corresponding to progress through the task. This allows the variable length problems to be mapped to a common slot representation. Here 3 additional slots are shown between slot 1 to 2 (as well between slot 2 to 1) corresponding to 25%, 50%, and 75% progress. This models says that RNN neurons will be tuned to ‘task-progress’ since a preferred observation, e.g., a neuron will fire when ‘3’ was 125% progress lengths ago. **D)** The RNN learns task-progress slot cells, i.e., cells that fire at a consistent task-progress post an observation, as predicted by our model (panel C). Top: RNN neuron that fires at 0% after observation ‘7’. Left/Middle/Right: Average neural firing in all problems without a ‘7’ observation / problems with observation ‘7’ in ‘A’ position / problems with observation ‘7’ in ‘B’ position. Bottom: RNN neuron that fires at 175% after observation 5. Left/Middle/Right: Average neural firing in all problems without a ‘5’ observation / problems with observation ‘5’ in ‘A’ position / problems with observation ‘7’ in ‘5’ position. **E-G)** Same as A,B,D but for a 4-ISR task where delays between observations can be different for different problems (but still fixed within each problem; example delays of 3, 0, 1, and 4 steps shown in schematic; delays sampled from {3, 4, 5}).

**Figure 5.**
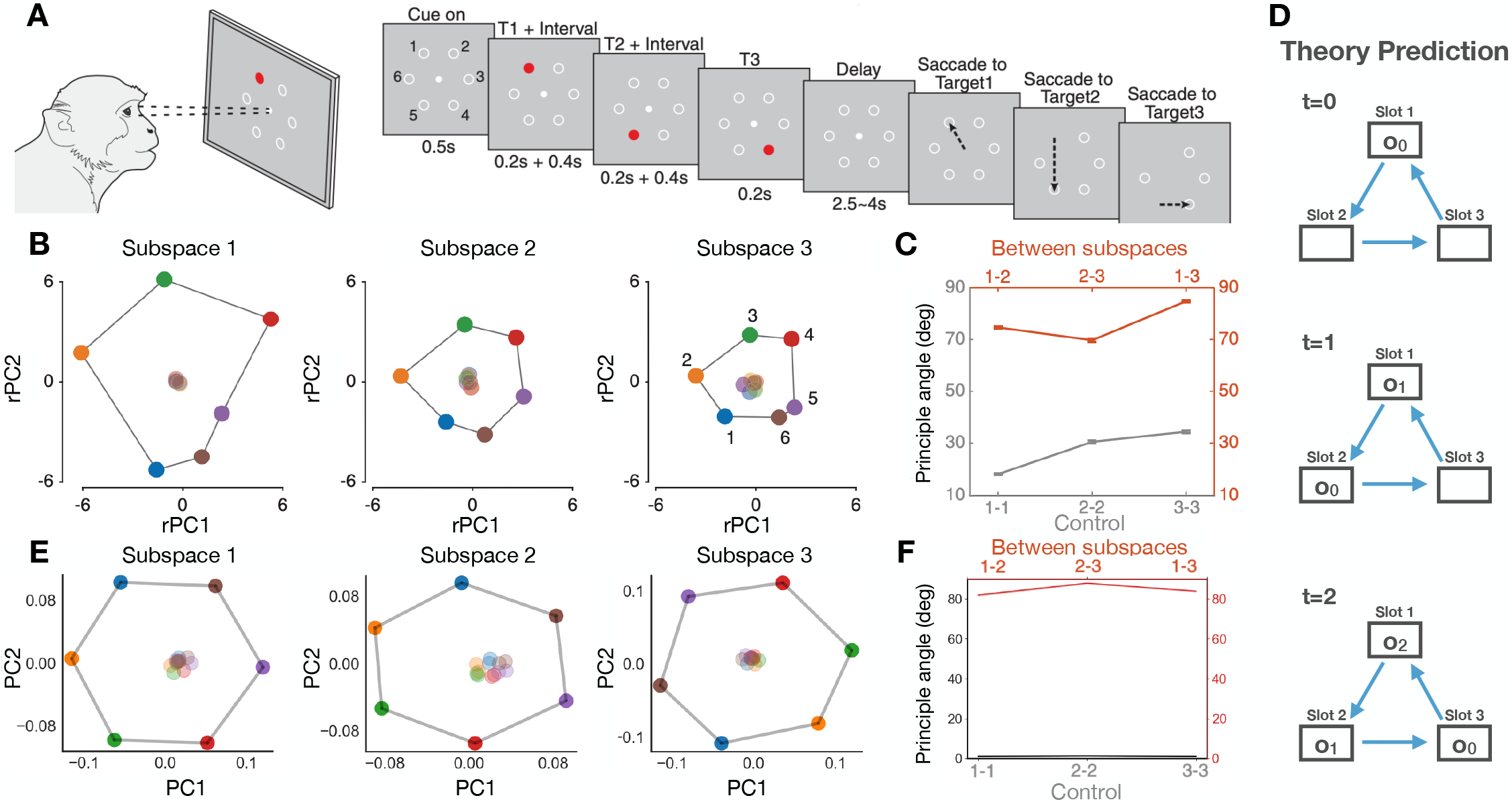
Explaining PFC representations in an ISR task as activity slots. **A)** Xie et al.^29^, recorded PFC neurons while monkeys performed an ISR like task. Monkeys were presented, on a screen, three (out of six) spatial positions in sequence, then, after a delay, tasked with repeating the sequence (via saccades). **B)** During the delay, PFC recordings revealed observation coding in three neural subspaces; one for each ordered position. Darker dots are projections of the *i*^th^ observations in the *i*^th^ subspace, lighter dots are projections of *j*^th^ observations in the *i*^th^ subspace (*j* ≠ *i*). Lighter dots have close to zero projection, thus suggesting the subspaces are orthogonal, which **C)** is confirmed in analysis. **D)** For this ISR task, the activity slots model adds the current observation memory to slot 1, and rotates the contents of the slots unidirectionally, so that after observing all observations, slot *i* contains the observation *i* − 1 timesteps ago. In this schematic, the readout slot is also slot 1. **E-F)** Training a RegularRNN on the same task recovered the same three orthogonal subspaces, and is consistent with an activity slot model. A-C) reproduced from Xie et al.^29^.

Indeed, after training WM models on these tasks, we are able to decode progress, as well as progress-velocity (Figures 4A,E right). Furthermore, we observe progress tuning at the single neuron level (Figure 4B,F), as well as progress-velocity tuned neurons (Figures 15A, 16A). This means the network internally computed progress-velocity, on the basis of sparse inputs, to overcome the different delays. The question remains; how does this relate to activity slots?

**Figure 6.**
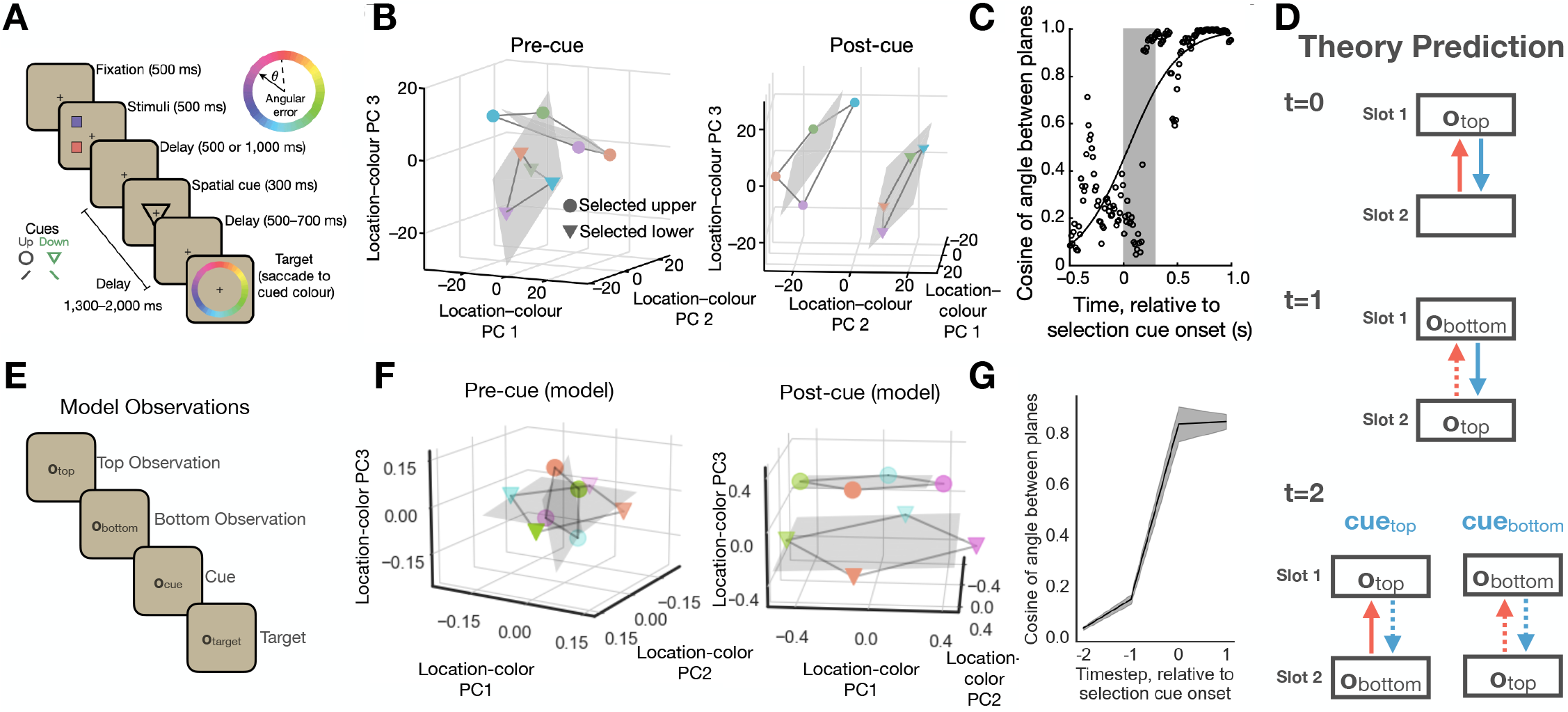
Explaining PFC representations in a cue-dependent memory retrieval task as velocity controlled activity slots. Panichello & Buschman^33^ recorded PFC neurons while monkeys were cued to recall one of two colours previously presented at the top and bottom of a screen. **B-C)** Before the cue, PFC neurons represented two orthogonal subspaces - one for the top colour and one for the bottom colour. After the cue the subspaces became parallel. **D)** For this task, the activity slots model adds observation memories to slot 1, and copies and shifts the contents of slots according to velocity control signals (direction of shift denoted by solid as opposed to dashed arrow). In this schematic, the readout slot is also slot 1. **E)** We modelled this task by sequentially presenting the top colour, the bottom colour, then the cue, and trained the RNN to predict the target colour. **F-G)** A trained RegularRNN recovered the same orthogonal subspaces, which became parallel after the cue. This is consistent with the activity slot model. A-C) reproduced from Panichello et al.^33^

**Figure 7.**
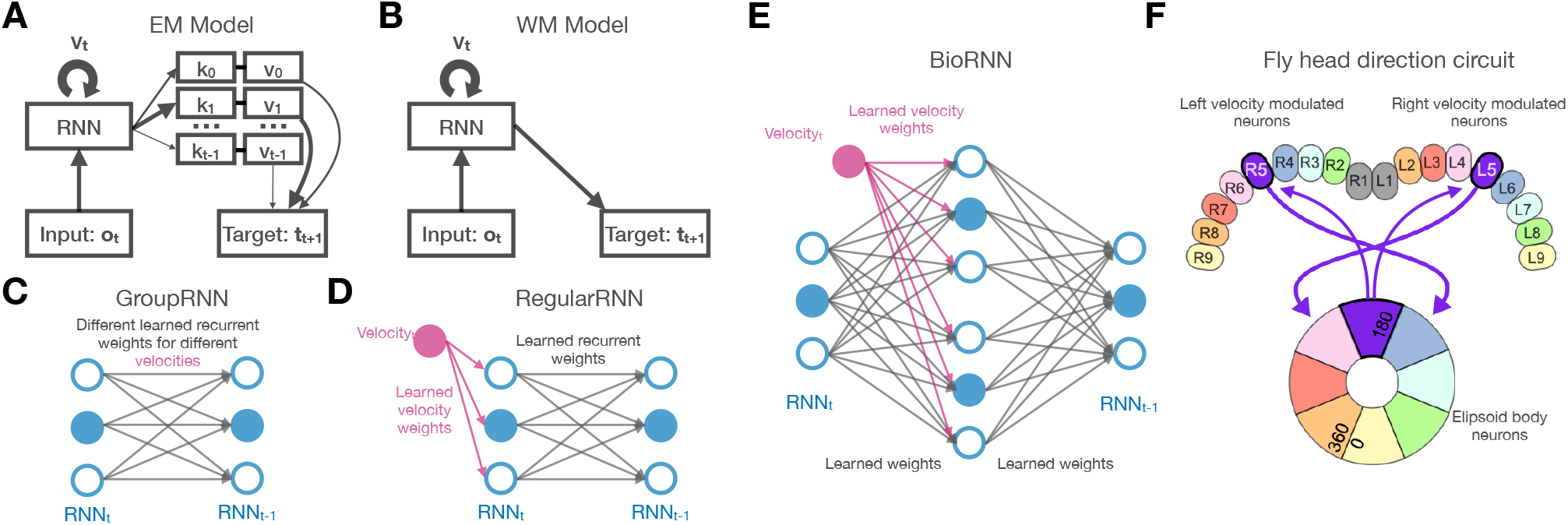
Model variants. **A)** The structure of the EM model. An RNN stores past memories using keys and values, like a transformer model which is related to Hopfield networks. Predictions are made by the RNN indexing memories. **B)** The structure of the WM model is a standard RNN setup. **C)** Like our theory we use an RNN with learnable velocity dependent matrices (GroupRNN); **D)** We also use a conventional RNN which receives velocity as input (RegularRNN). **E)** The BioRNN variant integrates velocity in an intermediate step between transition, much like the **F)** Head direction circuit discovered in the fly brain^47^. (E) is adapted from^61^.

**Figure 8.**
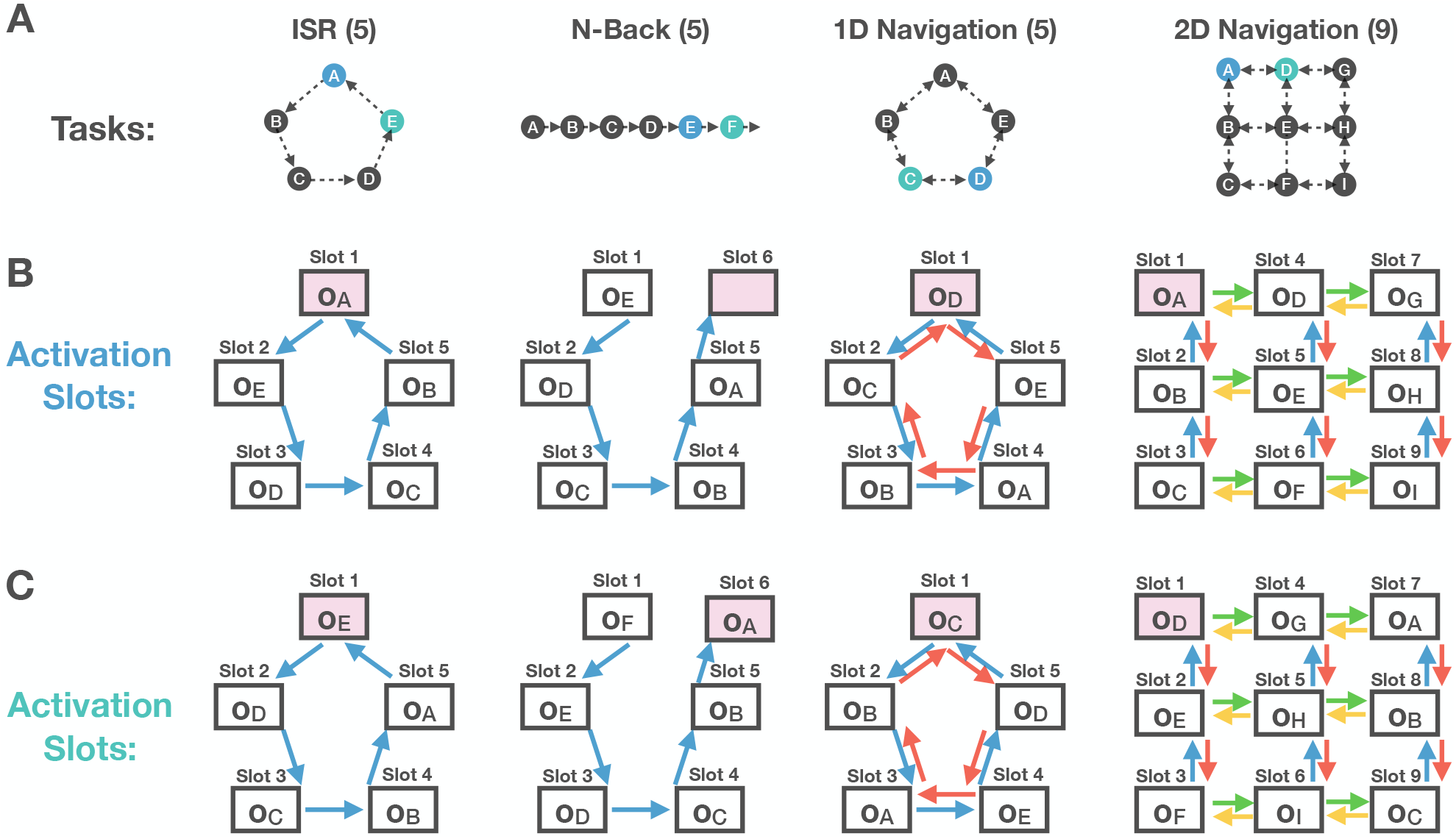
Tasks, and model predictions of activity slots. **A)** The structure of the four tasks. ISR is the same as unidirectional walks on a loop (5-loop shown). N-Back is unidirectional walks on a line (5-Back task shown). 1D navigation is a random walk on a loop (5-loop shown. 2D navigation is a random walk on a 2D torus (3-by-3 torus shown). **B)** The activity slot structure mimics the underlying structure of the task. Purple slots are the readout slots. Observations shown in slots, for the blue position above. The N-Back task nearly has a loop structure as observations more than N steps ago are forgotten. **C)** Observations now shown in slots, for the green position above. Note how the observations are copied and shifted between activity slots.

**Figure 9.**
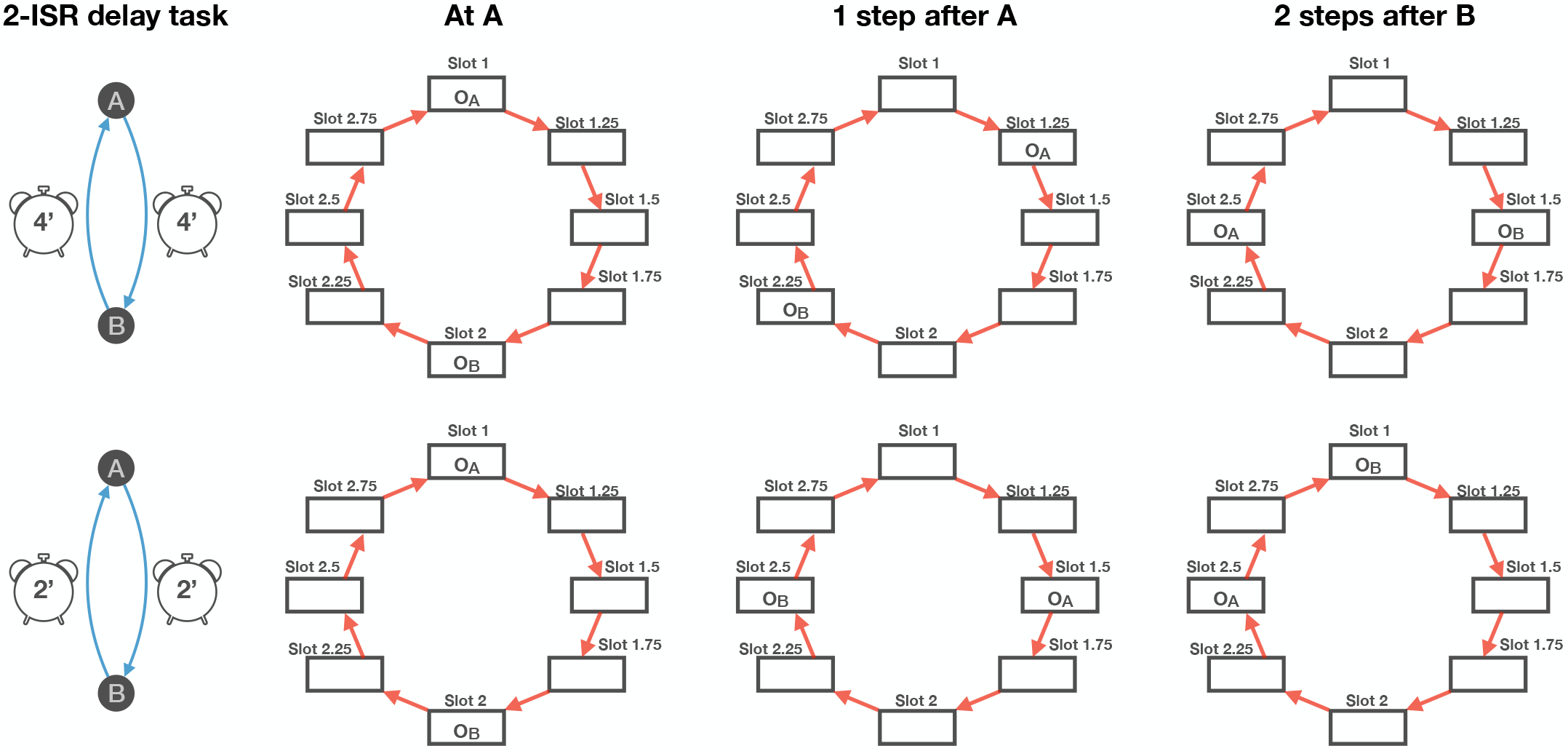
Task-progress slot prediction for 2-ISR delay task. We show two example 2-ISR delay tasks (top/bottom). The top task has delays of 4 steps, while the bottom has delays of 2 steps. For each task, we show task-progress slot predictions at three different time instances. We see differential activity between the two tasks, since the delay periods differ and so observations move round the task-progress slots at different rates.

**Figure 10.**
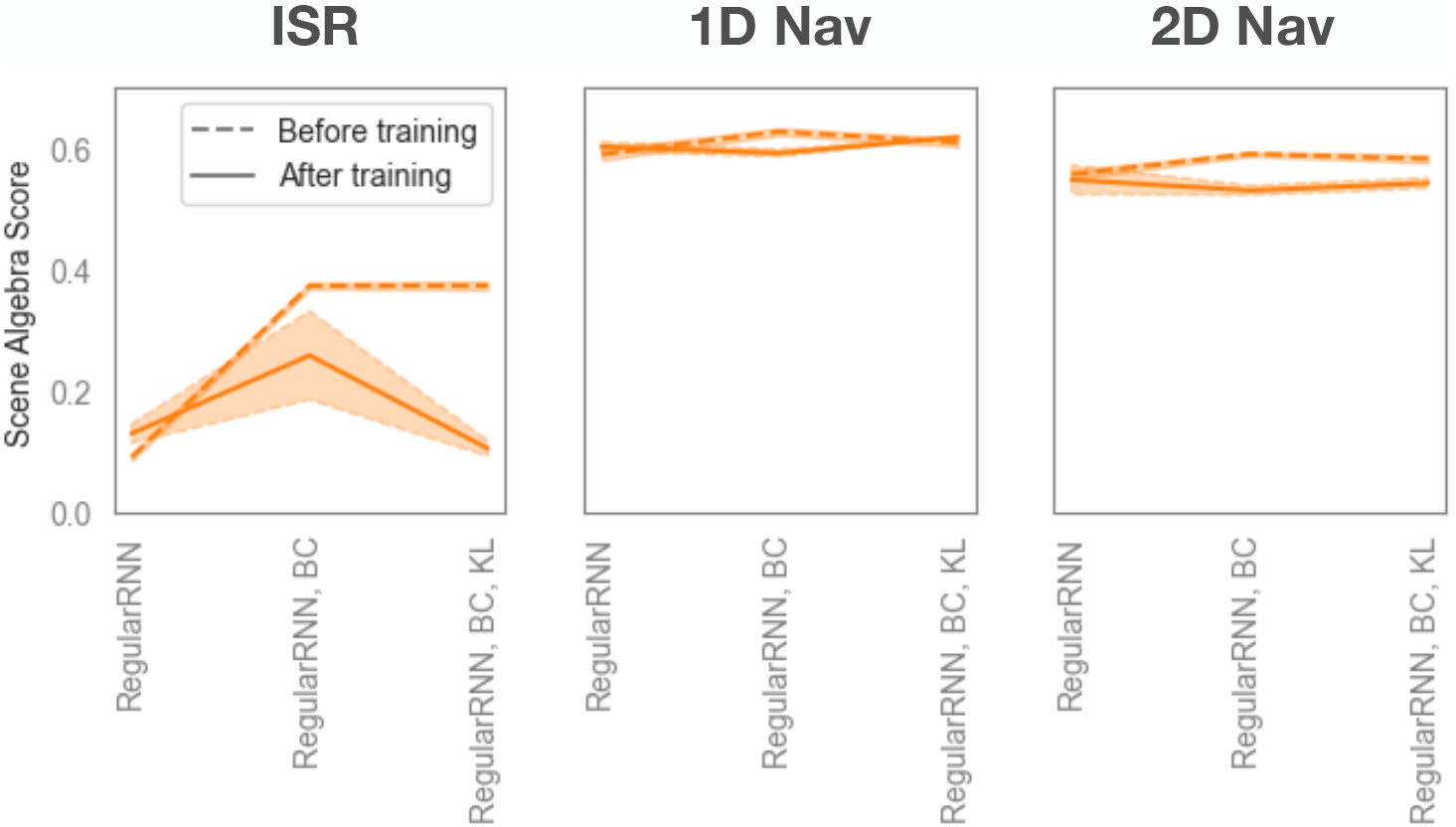
Slot algebra for RNN receiving velocity input during ‘path integration’: The same slot algebra analysis as Figure 3C, but with a RNN where the velocity signal of the upcoming transition is only provided to the path integration step, i.e. ***r***_*t*_ = ***W*** _*rec*_***h***_*t*−1_ + *B****v***_*t*_ (***B*** is a learnable matrix), and the external input, ***i***_*t*_, is just ***o***_*t*_. Slot algebra scores are high for tasks that involve variable velocity signals (1D Nav and 2D Nav) suggesting that the learned internal representation is not invariant to trajectory.

#### Controlling activity slots with progress-velocity

We posit that activity slots get structured by both task and progress (examples for the 2-ISR delay task shown in Figures 4C, 9), in that the slots are organised by task-progress on a loop. This means a slot will now code for 25% since observation, or 150% since observation (up to 200/400% since for the 2/4-ISR delay task). We refer to this as ‘task-progress’ structured (Figure 4C). The contents of these slots are copied and shifted to subsequent slots by the progress-velocity signal^iv^. Indeed, in trained WM models, we are able to decode task-progress activity slots (Figure 4A,E right), and we observe ‘task-progress slot’ neurons in Figure 4D,G; these neurons always fire at the same task progress after a preferred observation (e.g., 375% after observation 5; Figure 4G bottom). Further cellular examples are shown in Figures 15, 16. It is of particular note that both ‘progress’ and ‘task-progress slot’ neurons were recently recorded in rodent mPFC^30^, exactly as our model predicts.

While representing progress and progress-velocity seems an esoteric choice of representation, it is not clear how activity slot representations could solve this task without representing progress-velocity. For example, it is not possible to have an activity slot for each step of the delay period as this cannot be placed in a consistent loop structure different problem realisations of a task. In general, when behaviour (e.g., navigating to goals or waiting for delays) is structured in different coordinates to the underlying space where actions are taken (e.g., physical space or a loop), a coordinate transformation is required to match the two spaces. This is exactly what the progress, and progress-velocity, neurons are doing: transforming from spatial coordinates to task coordinates.

#### PFC activity slots in a task with constant velocity

Our activity slot theory also accounts for PFC representations in standard sequence memory tasks where only a constant velocity is required. For example, Xie et al.^29^ recorded PFC neurons in an ISR task. Here, monkeys were shown a sequence of three spatial positions on a screen (out of 6 positions), and, were trained to saccade to the same objects in order (Figure 5A). Importantly, the monkeys were trained on many such sequences (i.e., just like our ISR task). Neural activity in the delay period decomposed into three successive orthogonal subspaces. Each of the 6 possible spatial saccade locations could be represented in each of the 3 subspaces, with the first/second/third saccade location encoded in the first/second/third subspace (Figure 5B-C). These 3 sequential subspaces therefore directly correspond to our activity slots, with a different slot for the first, second and third ordinal points in the sequence (Figure 5D bottom is after delay period). Indeed, a WM RegularRNN on trained the task, exhibits exactly the same subspace decomposition (Figure 5E-F).

#### Controlling PFC activity slots in a cue-dependent memory retrieval task

So far we have considered either external or internally generated velocity signals. But sensory observations themselves can also serve as velocity signals. Here we show that sensory cues can be understood as velocity signals controlling PFC activity slots, in a particular study of cue-dependent memory retrieval^33^. In this study, Panichello & Buschman^33^ recorded PFC neurons in monkeys performing a working memory task where two colours are presented on a screen, one at the top and one at the bottom. Then, after a delay with a blank screen, a cue informs the monkey whether to report the colour of the top or bottom colour (Figure 6A). Importantly, each problem realisation is a random combination of two colours and cue. Neural activity in the delay period decomposed into two orthogonal subspaces, one for the top colour and one for the bottom (Figure 6B, left). Thus, each colour can be represented in one of two subspaces, depending on whether it was presented at the top or bottom. After the cue, the subspaces rotate so that top (when correct) and bottom (when correct) subspaces are parallel to one another (Figure 6B, right; Figure 6C). This is easily understood with our theory (Figure 6D); the initial two subspaces are two different activity slots, one for the top colour and one for the bottom colour. Since only one slot can be readout from, the cue can be understood as a velocity signal that controls how slot contents are shifted to the readout slot. Because either the top or bottom colour is moved (controlled by the cue (e.g. velocity) signal) to the same readout slot on correct trials, the subspaces become parallel to one another after the cue on correct trials (Figure 6D, bottom). Indeed, a WM RegularRNN trained on the task, exhibits exactly the same subspace decomposition and dynamics (Figure 6E-G).

## Discussion

In this work, we derived a formal relationship between the algorithms and representations of episodic and working sequence memory, and empirically validated theoretical predictions that EM utilises more abstract position-like representations while WM uses activity slot representations with a particular and systematic conjunctive coding of observations and (counterfactual) actions. Furthermore, we show that, like the HPC EM system, the PFC WM system can be updated and controlled by velocity signals, which can be flexibly computed from sparse input data. We used our theory of controllable activity slots to show that recordings from several different PFC studies - from serial recall to cue dependent recall and spatial working memory - are all explained in a unified framework of activity slots controlled by velocity signals.

Intriguingly, our activity slot theory may also explain PFC data from non-explicit working memory tasks. For example, human mPFC (fMRI) activity obeys rules of ‘scene algebra’ when humans perform scene construction tasks involving one object above another^52^, i.e., 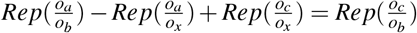, where 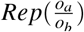 denotes the representation of object *a* above object *b*. This suggests that the objects are stored in different neural subspaces depending on their spatial location, analogous to our theory of activity slots which organise objects depending on their (e.g. spatial) relationships. This is perhaps no surprise since both our and their task require a compositional understanding (objects can be in any sequence or spatial configuration respectively). We anticipate a slot based understanding will offer further insights into representations in frontal cortex, beyond just working memory. For example, activity slots could represent sub-goals for sequential goal directed planning^30^. Each slot would hold a sub-goal, with the sub-goals cycled forward when the previous sub-goal is achieved. This could offer an explanation as to why the same brain region involved in working memory, is also implicated in value based decision making and goal directed planning^31,53^, and why there are goal predictive representations in PFC (recall neurons in activity slots conjunctively encode different degrees of distance/progress to various observations/goals). In this vein, it is of note that slot based models are used in machine learning in diverse settings from natural language^6^, to planning^54^ and scene construction^55,56^.

We related the two algorithms to HPC and PFC, however the key distinction between the algorithms is whether memories are stored in synaptic strengths or neural activity patterns. It is plausible that PFC could utilise short term plasticity to store memories in synapses as well as neurons. Indeed, recent modelling work suggests short term plasticity may better explain PFC activity as compared to recurrent dynamics^57^. Conversely, because of the intimate connectivity between entorhinal cortex and hippocampus (compared to the more distant connectivity between PFC and hippocampus), it makes sense for entorhinal cortex to control hippocampal synaptic memories using position-like representations as opposed to activity slots.

We do not claim to have solved the representations of RNNs on all working memory tasks. Indeed there are WM tasks in which activity slots are not learned (e.g., Figure 14A-B). However, we suggest that activity slots are a general solution to sequence working memory tasks, but in some circumstances there may be other bespoke RNN solutions that get learned instead. Lastly, we note that our formalism, as well as the GroupRNN, is a state-space model (these models have gained considerable attention recently^42^), thus, on the machine learning side, our work offers a relationship between the learned representations of state-space models, RNNs, and transformer neural networks.

**Figure 11.**
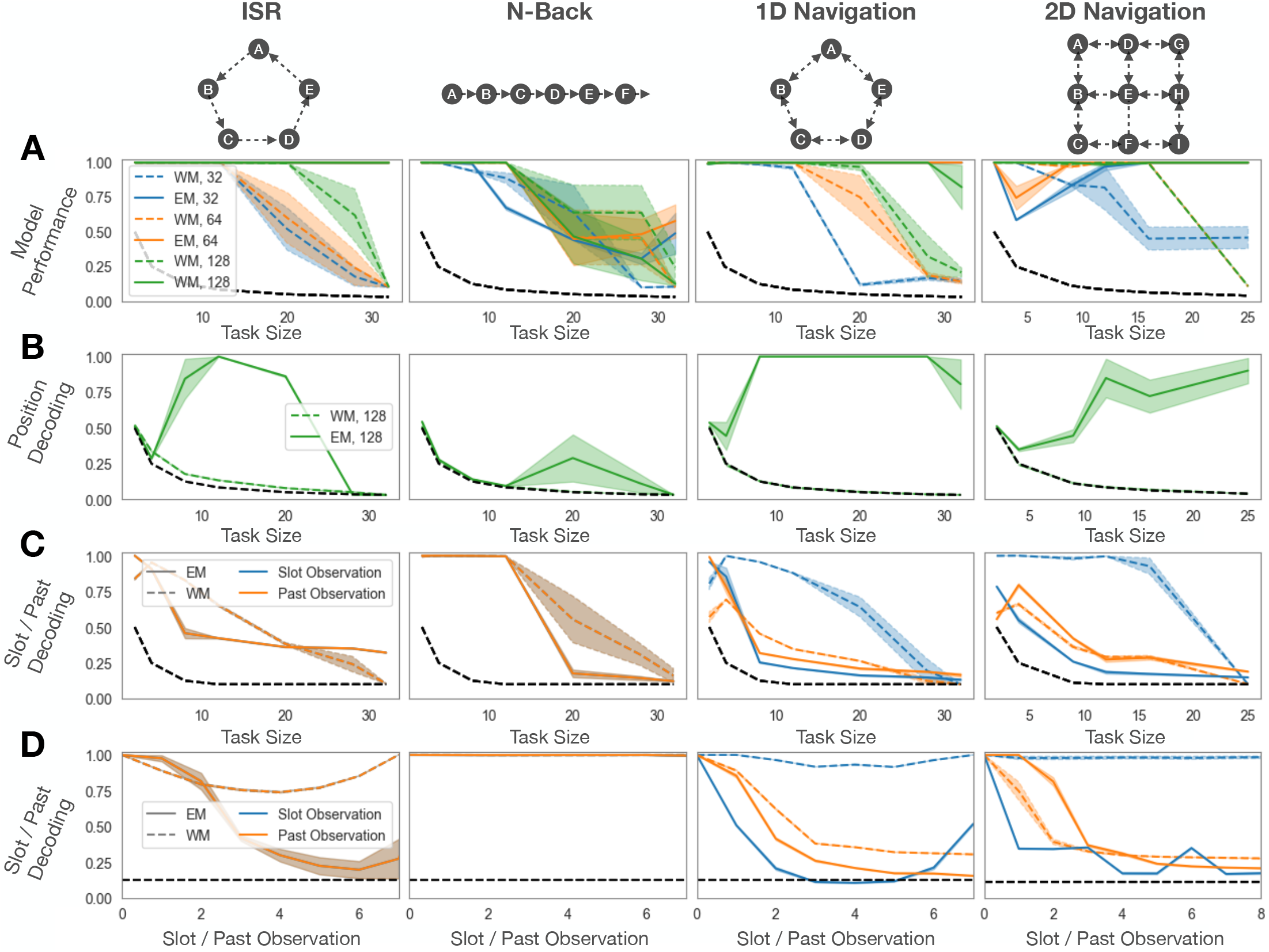
GroupRNN: EM models scale better and learn position representations, while WM models learn activity slots. For each combination of task, task size, model type, and RNN size (32, 64, 128), we train five randomly initialised models. Results are mean *±* standard error. **A)** Performance of the EM (solid) and WM (dashed) models on the four tasks. EM models consistently perform better that WM as task size increases. Larger sized RNN consistently perform better for both EM and WM models. All subsequent panels are for RNNs with size 128. **B)** Position decoding. EM models, but not WM models, learn position like representations. **C)** Average decoding of ‘slot-sequences’ (which observations should be in each slot at each timestep according to our theory) and ‘past-sequences’ (what observation occurred *i* steps ago, up to the total number of slot). For ISR and N-Back tasks, slot- and past-sequence decoding is identical since velocity is constant (+1) thus the *i*^th^ past observation will be held in the *i*^th^ slot. Otherwise, slot decoding is consistently higher than past-sequence decoding for WM models, but not EM models. **D)** Slot- and past-sequence decoding for individual slots or past timesteps (on task with size 8 for first 3 tasks, and 9 for 2D Navigation). Slot decoding is consistently higher than past-sequence decoding for WM models, but not EM models. We note that on the ISR task, performance is high but position decoding is low for large tasks sizes. We posit this is due to an additional solution only available in this task: a Laplace transform. We further note that for the N-back task, the position (or slot) we decode is not the veridical position (or slot) since positions (or slots) never get repeated. Instead, we decode position (or slot) as if it were cycling on a N-loop. Thus it is the structure of the input-output map that gets learned, rather than the structure of the input.

**Figure 12.**
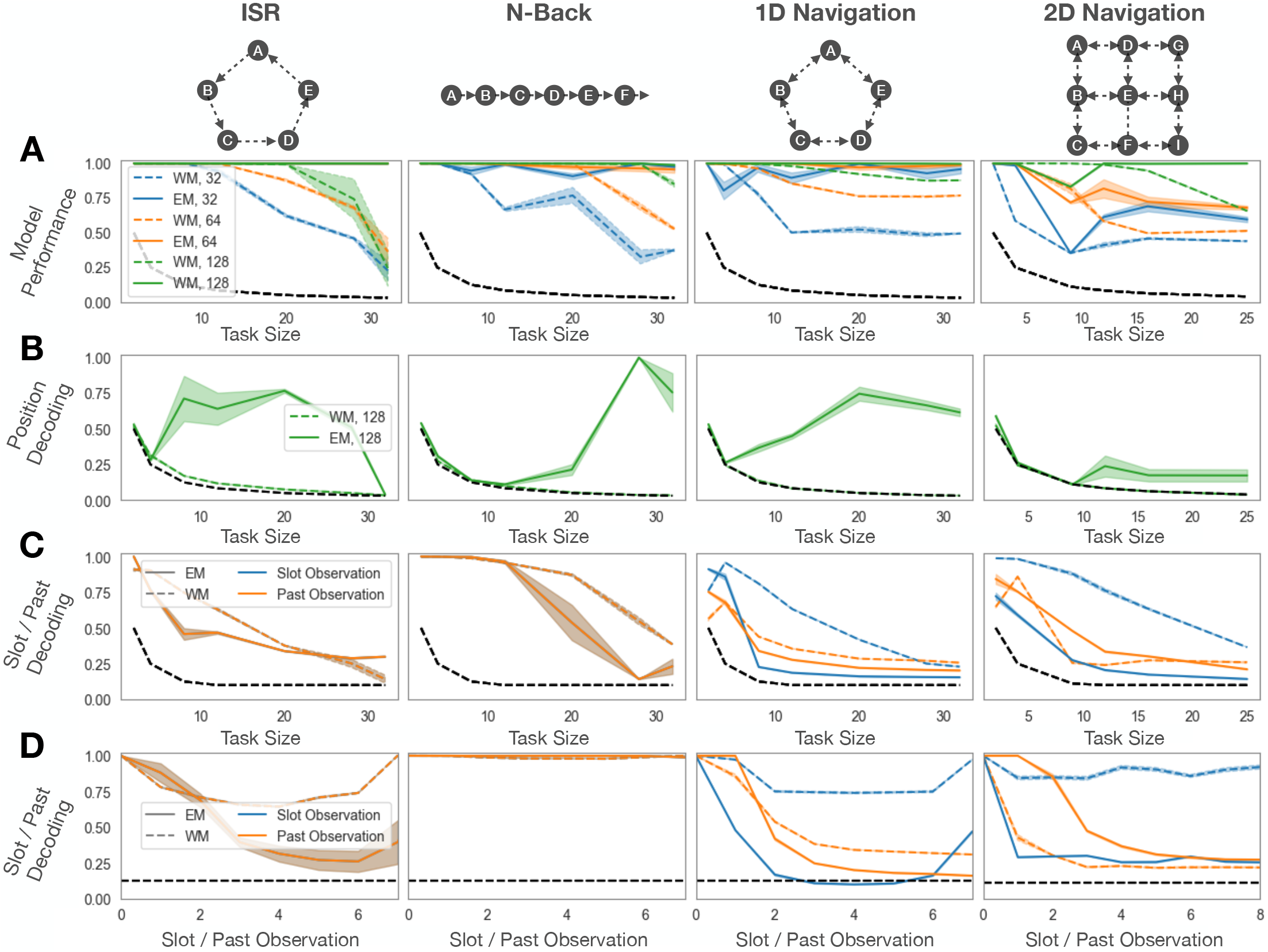
RegularRNN: EM models scale better and learn position representations, while WM models learn activity slots. Same as Figure 11 but for the RegularRNN

**Figure 13.**
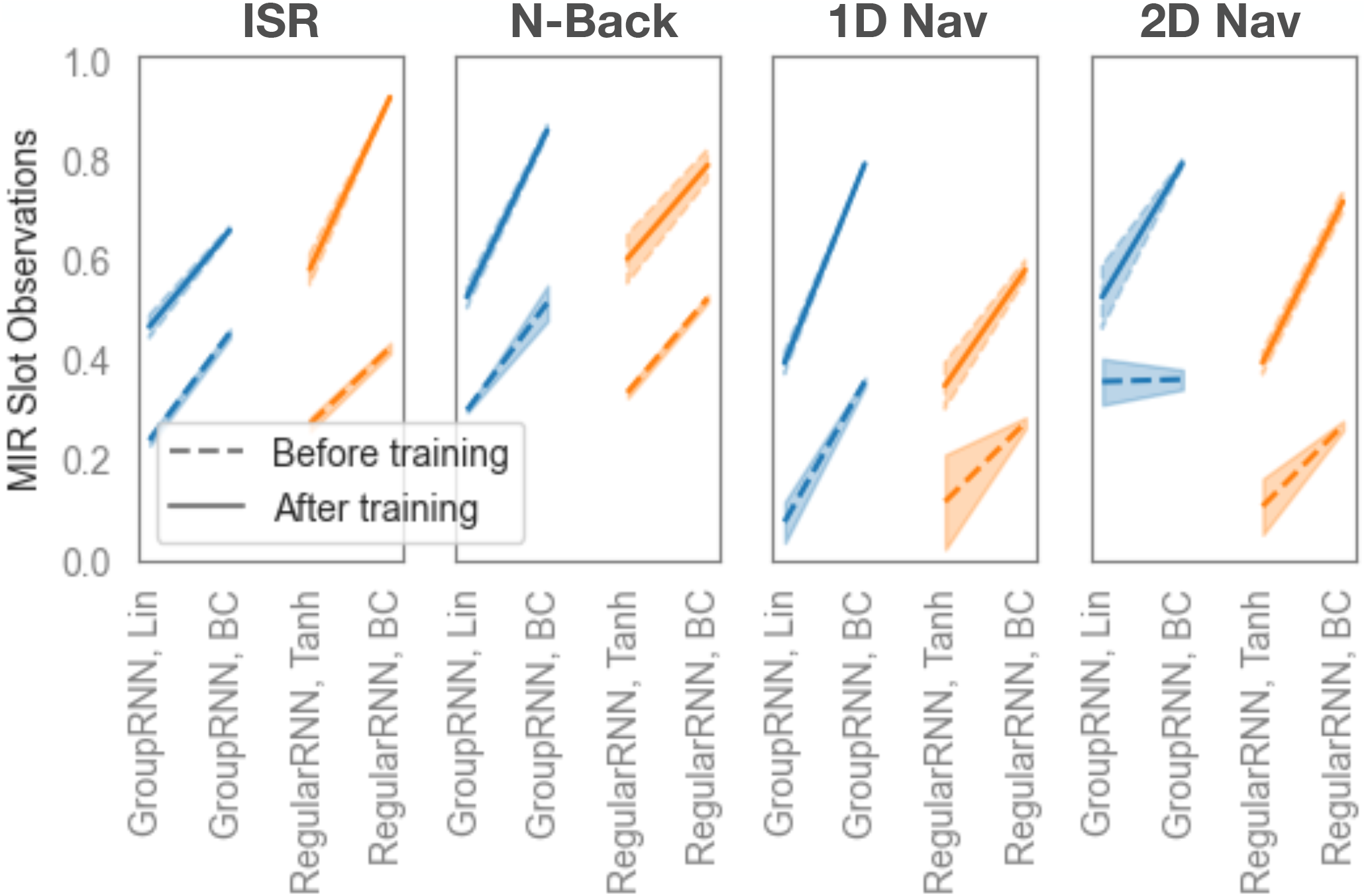
WM models display single neuron coding of slots when additional constraints (BC) are used. The Mutual Information Ratio (MIR) measures the extent of single neuron coding of slots. MIR rises from the beginning of training (dashed) to the end (solid), particularly for models with additional constraints.

**Figure 14.**
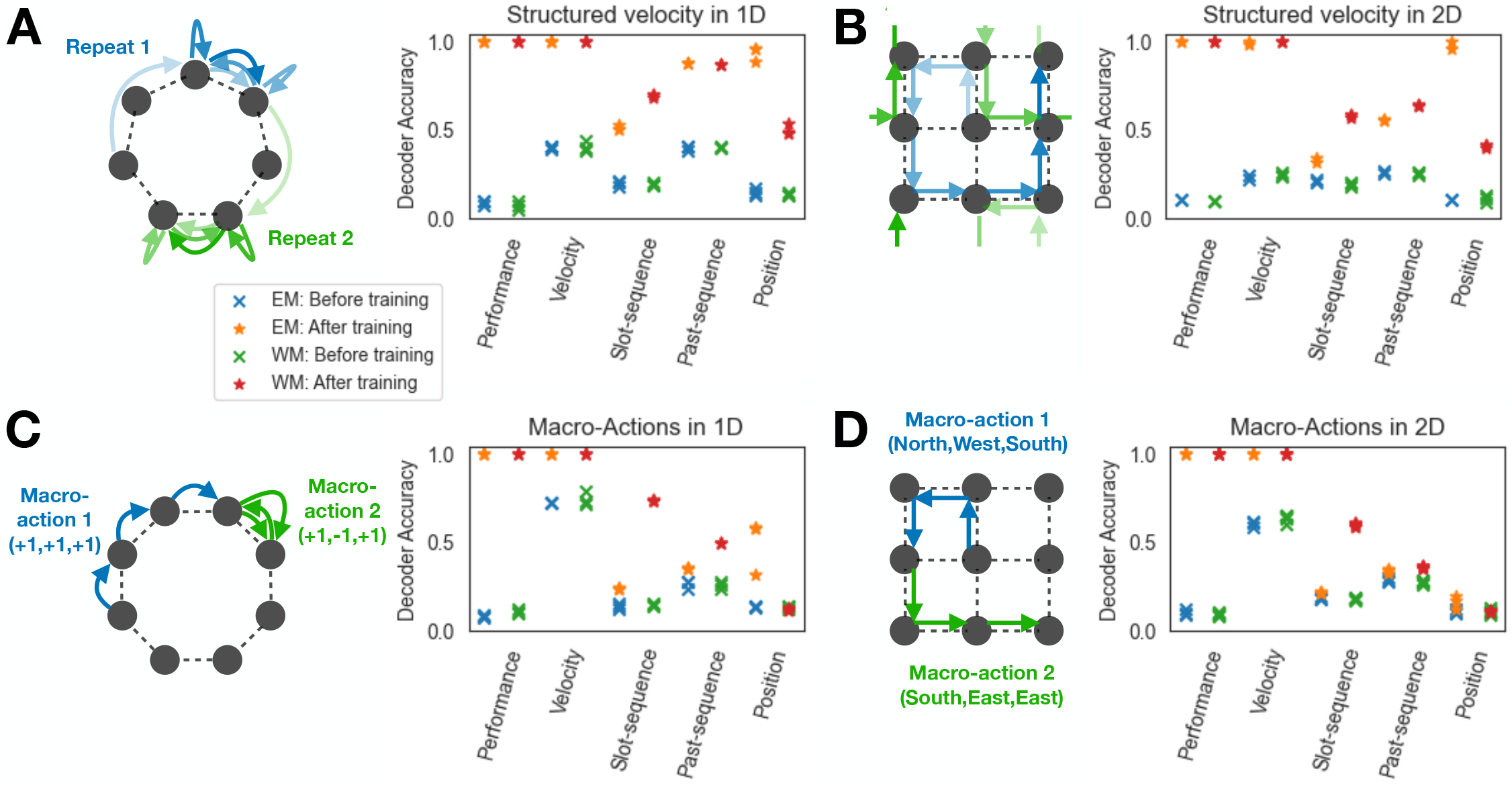
Activity slots are not learned when velocities are non-random sequences. **A)** Left: On a 7-loop task (1D navigation), but where velocity cycles a fixed schedule of {+2, +1, 0, − 1, 0, +1, +2, +1, 0, … } Schematic shows 1st / 2^nd^ cycle of fixed velocity schedule shown in blue / green. Right: Each point corresponds to a randomly initialised model before (cross) and after (star) training; three random initialised models shown. Velocity can be decoded in both EM and WM models. WMmodels, however, do **not** learn activity slot representations: past-sequence decoding is higher to slot-sequence decoding. We note that EM models still learn a position representation. **B)** Same as A) but for a fixed velocity schedule ({ `North`, `West`, `South`, `South`, `East`, `East`, `North`, `North}`) on a 3-by-3 torus. **C)** Left: Taking macro-actions on a 8-loop task (1D navigation). The input ‘velocity’ signal is each macro action, which can be up to 4 sub-actions long. Schematic shows 2 macro actions in blue / green. Right: Each point corresponds to a randomly initialised model before (cross) and after (star) training; three random initialised models shown. The model learns to represent the sub-actions in both EM and WM models (decoder accuracy shown for final sub-action). WMmodels **do** learn an activity slot representation, while EM models learn a position representation. **D)** Same as B) but in a 2D navigation task on a 3-by-3 torus. These results suggest that activity slots are not learned when velocity signals are fully structured (like the prescribed velocity sequences in A and B), but when randomness is introduced (like different random macro-actions, macro-actions are in-between structured sequences and fully random actions) activity slots are learned.

**Figure 15.**
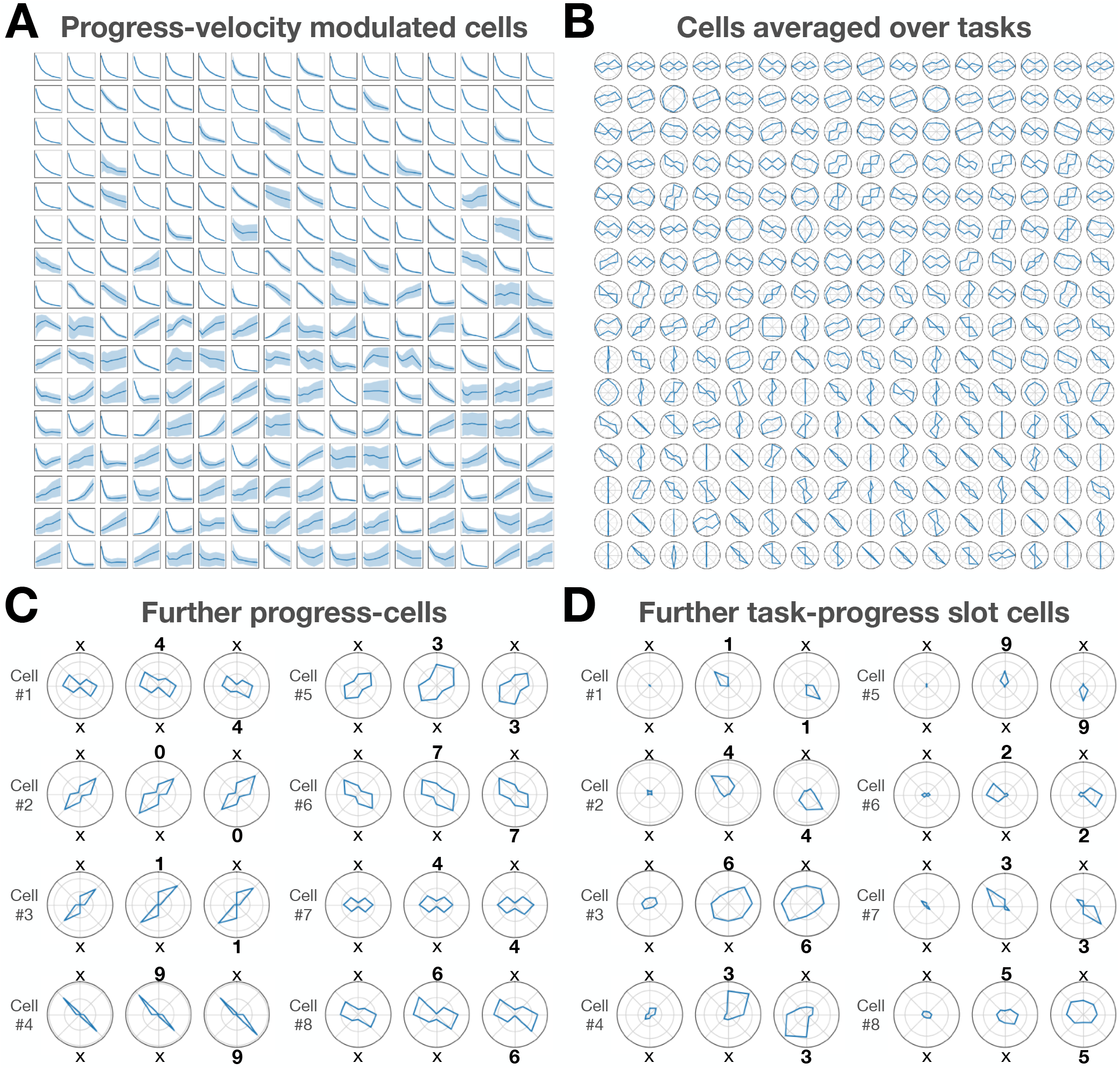
Further RNN cells representations for the 2-ISR delay task in Figure 4. **A)** RNN cells are modulated by progress-velocity (showing 256 of 512 cells with highest activity). Each subplot is a cell. The x-axis is the ‘true’ progress-velocity (i.e., 1 divided by the current delay length). The y-axis is the average firing of that cell for that progress-velocity. **B)** The average firing of RNN cells across many tasks (we rescale activity around the 2-ISR delay loop into a common length, i.e., task-progress; we show 256 of 512 cells with highest activity). The average firing across tasks for most cells is bimodal. This is consistent with both progress (bi-modal) and task-progress slot cells (uni-modal but rotated across tasks). **C-D)** In these plots, we have extracted the ‘preferred’ observation for that particular cell (the observation that, when present in a task, leads to the cell having the highest activity), and plotted the cell’s activity when the preferred observation is at different positions on the 2-loop. Left: Observation not present. Middle/Right: Observation in A/B position. **C)** Example progress cells. Progress cells respond the same, even for tasks without the preferred observation (left of each subplot; because progress cells do not have a preferred observation). Progress cells are bimodal as they track progress within the delay. **D)** Example task-progress slot cells. These cells do have a preferred observation; they are inactive when that observation is not present in the task (left of each subplot). They fire at a particular lag after their preferred observation, and so these cells are not bimodal.

**Figure 16.**
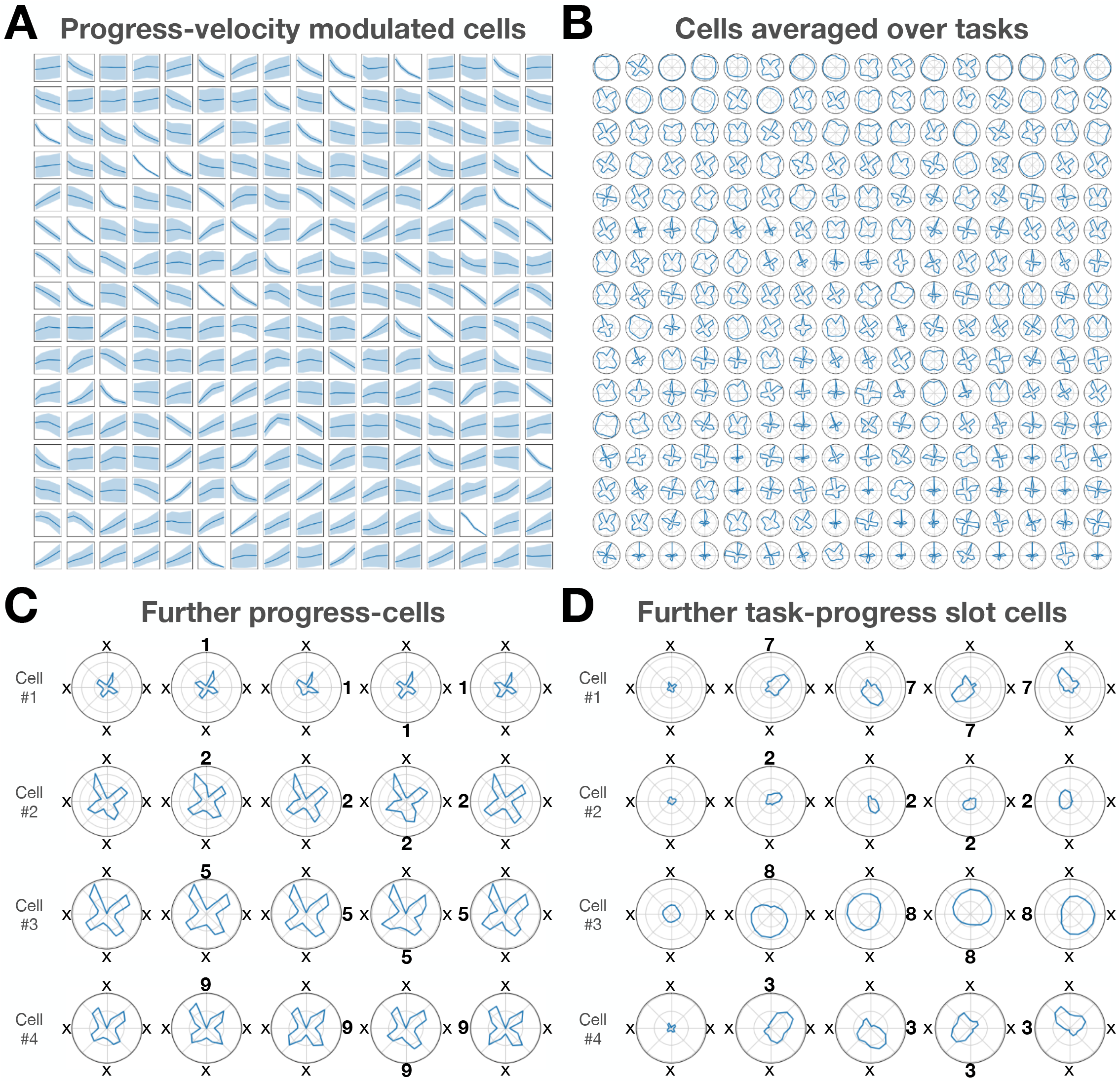
Further RNN neural representations for the 4-ISR delay task in Figure 4 A-D) Same as Figure 15, but for the 4-ISR delay task.

The hippocampal and frontal systems are thought to be essential in mediating higher order cognition, with patients exhibiting profound deficits in behavioural flexibility if these structures are damaged^10,11^. While these systems have been the subject of intense study over the last several decades - both experimentally and theoretically - the systems have been largely treated independently and their respective cellular representations are hard to reconcile. Our work brings together these systems by showing that the algorithms and representations of frontal working memory and temporal episodic memory are two sides of the same coin.

## A Appendix

### A.1 Full Theory

#### Task reminder

Formally, for each task, we consider a dataset *𝒟* = {*𝒟*_0_, *𝒟*_1_,*…, 𝒟*_*N*_ }, where each *𝒟*_*i*_ is a sequence consisting of vectors of sensory observations, ***o***, of dimension *d*_*o*_ (termed 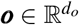), sensory targets, 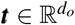, and (allocentric) velocities, 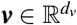, i.e., *𝒟*_*i*_ = {(***o***_0_, ***t***_0_, ***v***_0_), (***o***_1_, ***t***_1_, ***v***_1_),, (***o***_*K*_, ***t***_*K*_, ***v***_*K*_)} if the sequence is of length *K*. These velocities mimic real actions taken by agents, but in this case we provide them externally. Importantly, the underlying structure of the task prescribes how the velocities add up to cancel each other out (just like `North` + `East` + `South` + `West` = 0 defines 2D space), and when the velocities cancel each other out, the observation is identical to what it was previously (just like returning to the same position in 2D space). Concretely, there is an underlying latent variable, 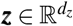, corresponding to ‘position’, and each position, ***z***, is associated with an observation, ***o***, and any two neighbouring positions are related by a velocity, ***v*** (example for a simple loop structure in Figure 1A left). For each *𝒟*_*i*_, while the observation-position pairing is random, the underlying structure is preserved (velocities add up and cancel in the same way). The aim of the task is to predict a target, ***t***, which is either an upcoming observation (i.e., what you will see after going `North`), or a past observation (i.e., what you saw 5 steps ago). Importantly, at time *t* the model predicts the target at time *t* + 1 - one-step prediction. While the task formalism (and subsequent model formalism) is general to tasks with discrete and continuous positions, we primarily consider the discrete setting here (with *n*_*p*_ total positions).

#### Solving these tasks with an EM model

Such tasks can be solved using EM systems that consist of RNNs equipped with an external memory. The external memory can be a Hopfield network^4^, a modern Hopfield network^35^, a transformer neural network^36^, or a differentiable neural dictionary^58^. All these methods are relatable to each other^36^, and crucially can all be viewed as storing memories in synaptic connections. Here, we present the differential neural dictionary / modern Hopfield network version (as they are general external memory devices). Additionally, we consider an RNN that only receives recurrent input and the velocity signal. Predictions are made via

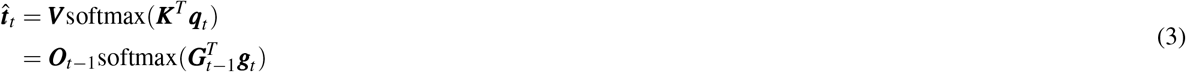

Where 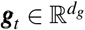 is the RNN state representation at time *t*, and ***O***_*t*− 1_ = [***o***_0_…***o***_*t*− 1_] and ***G***_*t*− 1_ = [***g***_0_…***g***_*t*− 1_] are stored memories, i.e., the memories bind together each ***g*** and ***o*** at each timestep^v^. We note that the targets are in the form of observations (upcoming/ those that have previously been seen) and so stored memories of observations will facilitate accurate prediction.

The particular choice of RNN does not matter for our purposes here - it just needs to accurately track position, ***z***. Nevertheless, the RNN state, ***g***_*t*_, will be a function of past velocities, ***v***, as it must integrate velocities to track position:

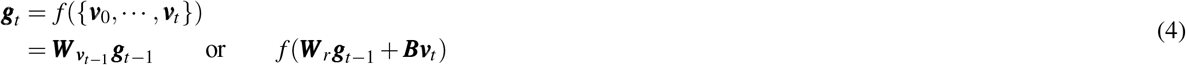

In the bottom line of the above equation, we present two common ways for integrating velocity in RNNs, the first^4^ uses velocity dependent matrices, ***W*** _***v***_, to update RNN state and is inspired from group theory (and can mathematically shown to produce grid cells in 2D space^40^), and the second^14^ is a classic RNN where ***W*** _*r*_ is a recurrent matrix and ***B*** is an matrix that maps velocities ***v*** to the RNN. Note ***v***_*t*_ is the velocity from ***z***_*t*_ → ***z***_*t*+1_, and so while the classic RNN takes ***v***_*t*_ as input at timestep *t*, it is really the velocity that was provided at *t* − 1 that gets used to update position (via the recurrent weights) at time *t* (that’s why in the velocity-dependent matrix case we use 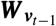 to get to ***g***_*t*_). This is just how classic RNNs are usually framed.

Regardless of the particular RNN, if ***g***_*t*_ learns to have the same structure as ***z*** (i.e., it represents position: ***g***_*t*_ = ***g***(***z***_*t*_)) then we can rewrite

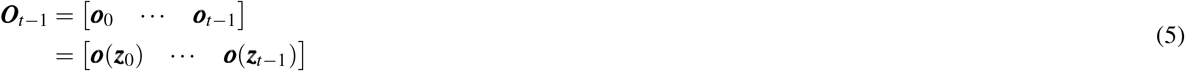

and

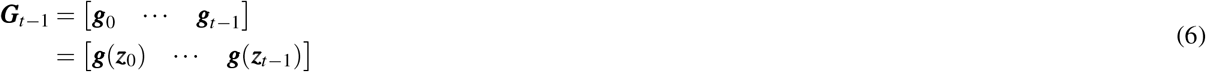

This says that the memories formed at each timestep (when at position ***z***_*t*_) are the *true* neural representation of position ***z***_*t*_: ***g***(***z***_*t*_) and the observation at position ***z***_*t*_: ***o***(***z***_*t*_). Thus the attention vector ***h***_*t*_ = softmax 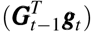will attend to the right memory at each timestep (i.e., memories that were paired to ***g* (*z*)** in the past) since ***g* (*z*)** will be more similar to itself, than to ***g* (*z***^*′*^)where ***z***^*′*^ is a different position.

#### Simplifying an optimal EM model

We consider a optimal EM model, and consider its representations. We assume that if an underlying ‘position’, ***z***, is visited more than once, the additional memory is not added, thus after all *n*_*p*_ ‘positions’, ***z***, have been visited 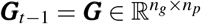 and 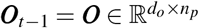, where *n*_*p*_ is the number of ‘positions’ for this task: we only store *n*_*p*_ memories. Thus, since a single memory get retrieved at any one time, the optimal attention vector, 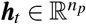 will be one-hot and with a single element active for each position^vi^. To make our lives easier, since attention is order invariant, we can relabel, and reorder, individual memories, not by the time, but by their ‘position’: 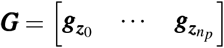 and 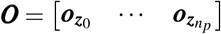. Since the one-hot attention vector, after a velocity ***v***, will change to another one-hot attention vector (coding for position ***z*** then position ***z*** + ***v***), the attention vector can be though of as representing a node on a graph. We call the attention vector at timestep *t* corresponding to position ***z***_*t*_ as ***h***(***z***_*t*_). Thus, the update to the attention vector is *functionally equivalent* to an attention vector being multiplied by an (velocity/action dependent) graph transition matrix, 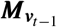, i.e., 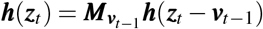. Thus (optimal) predictions can be rewritten as:

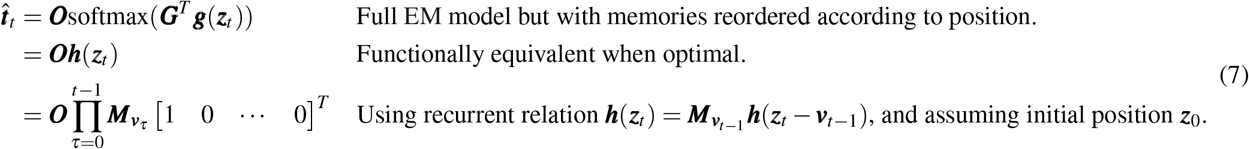

Expanding this equation out yields:

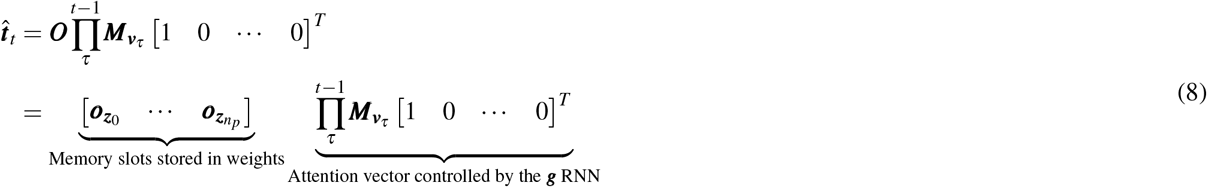

This equation describes the functional computations of an optimal EM model. We see that it reduces to an attention vector ***h*** that gets updates by velocity-dependent graph transition matrices. This attention vector then attends to memories stored in weights. Interestingly, this simplified formulation, can still be thought of as an RNN (the attention vector RNN) and an external memory (the memory slots). But we stress that this is a simplified formulation, as opposed to the actual model implementation (see Appendix A.2). Nevertheless it captures the computations of an optimal full EM model implementation.

#### Rearranging terms gives the WM solution

The above solution stores memories in synaptic connections. Here we show that reshaping and rearranging the above equation produces an alternative, but equivalent, solution where memories are stored in RNN activity rather than synaptic connections:

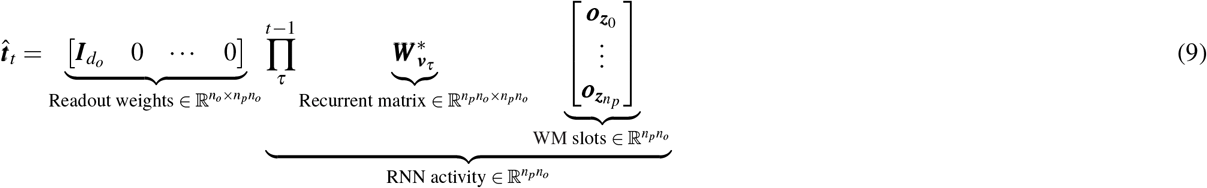

This equation performs the exact same computation as the previous equation, and its terms look very similar but there are some important differences (Table 1 for relationships between terms). For example, 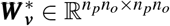 is like the previous velocity-dependent matrices, ***W*** _***v***_, but expanded (and transposed); where there was a 1 or a 0, there is now an identity matrix 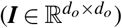 or a matrix of zeros 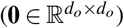. Intriguingly, what was a matrix of memories stored in weights, 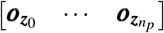 is now a vector of memories stored in RNN activity, 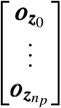.. Thus the neurons in the RNN can be though of as **activity slots** that store arbitrary memories in neural activity (Figure 1A right). Importantly, since the readout weights are now fixed, the contents of each slot must be copied and shifted to other slots via 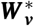, depending on the desired observation to readout. Adding a memory to a slot is simple and only requires feed-forward weights from observation to input slot (e.g., Figure 1A right). Thus, once learned, this model requires no synaptic plasticity. Lastly, similar to the EM solution, this WM solution works in any basis (not just the one presented above).

#### Separate attractor networks for computing velocities, and controlling slots

We now show that an RNN can consist of two components: a component for computing velocities, and a component for using velocities to control activity slots. A generic RNN updates is as follows:

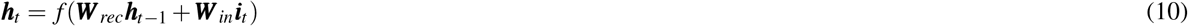

If we split ***h*** into two parts, ***h*** = [***h***^*s*^; ***h***^*v*^], where ***h***^*s*^ are the activity slots and ***h***^*v*^ are the neurons involved in computing velocity signals (we’re choosing a favourable basis where the two components are in separate neurons). Then we get the following:

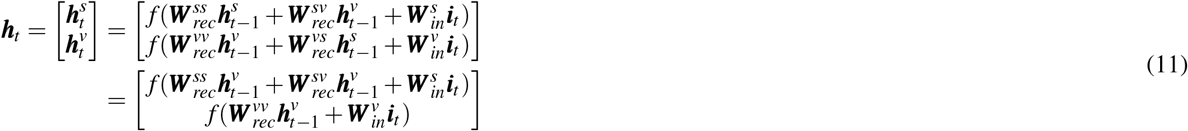

Where 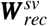 is the block of ***W***_*rec*_ that goes from *v* → *s* neurons etc, and in the last line we have assumed that information from slot neurons are not useful to velocity 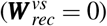. Thus this RNN consists of a velocity computing RNN, ***h***^*v*^, that computes velocities based on inputs and sends these velocities to the slot RNN, ***h***^*s*^, vis 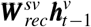.

### A.2 Model Details

We train models which are all variants on recurrent neural networks (RNNs), which differ in two key components (Figure 7). The two varying components are: 1) we use different RNN variants (varying from least to most biologically plausible), and 2) we use different methods of predictions from RNN activities for EM models (via fast Hebbian weights that change every timestep) and WM models (via slow weights learned by backpropagation).

#### RNN transitions

We use a gated RNN similar to a GRU^59^, where the external and recurrent inputs are gated: ***h***_*t*_ = *f* (***g***⊙ ***r***_*t*_ + (1− ***g***) ⊙ ***W*** _*in*_***i***_*t*_), where ***i***_*t*_ is the external input at timestep *t*, ***r***_*t*_ is the recurrent input, *f* (…) is an activation function, and 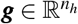 is an element-wise gate ***g*** = *σ* (***W*** _*α*,1_***r***_*t*_ + ***W*** _*α*,2_***W*** _*in*_***i***_*t*_) with the Sigmoid activation, *σ*. We compute the recurrent and external inputs in two different ways. First, like our theory, we use (learned) velocity-dependent matrices, 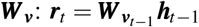, and the external input, ***i***_*t*_, is just ***o***_*t*_. We call this *GroupRNN* (Figure 7C). Second, we use a conventional gated RNN, with ***r***_*t*_ = ***W*** _*rec*_***h***_*t*−1_, and the external input, ***i***_*t*_, is ***o***_*t*_ and ***v***_*t*_ concatenated. We call this *RegularRNN* (Figure 7D). In Figure 3 we make use of another RNN inspired by the fly head-direction circuit (Figure 7E,F). In *BioRNN*, we have ***r***_*t*_ = ***W*** _2_ *f* (***W*** _1_***h***_*t*−1_ + ***v***_*t*−1_), where ***W*** _1_ projects ***h*** into a higher dimensional space. The external input, ***i***_*t*_, is just ***o***_*t*_.

#### Fast or slow readout weights

We use two readout methods to correspond to the two model classes in our theory. First, for the **WM** class: predictions are made by 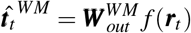, i.e. standard RNN prediction (Figure 7B). Second for the **EM** class: predictions are made using an external memory system related to a transformer / Hopfield network. In particular, 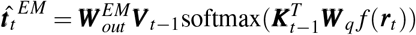, where the key and value matrix are sequentially updated i.e., ***K***_*t*−1_ = [***k***_0_…***k***_*t*−1_] and ***V*** _*t*−1_ = [***u***_0_…***u***_*t*−1_] (Figure 7A; we use ***u*** for ‘value’ as ***v*** is already used for velocity). At the beginning of each sequence, ***K***_*t*_ and ***V*** _*t*_ are reset to be empty.

#### Adding memories in EM models

Memories are simply added to a list, just like the differentiable neural dictionary^58^ or modern Hopfield network^35^. These memories consist of two parts - a key and a value (Figure 7A). The added key, as described above, at each timestep is ***k***_*t*_ = ***W*** _*k*_ ***h***_*t*_, while the added value is 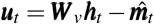, where 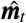 is the retrieved memory at that timestep 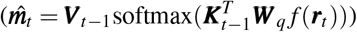. We do this, so that repeat copies of memories are not added.

Since the list of memories grows in time, we add an additional *β* scaling term in the softmax: 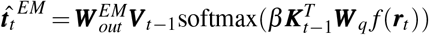. We do this as the normalisation term in the softmax sums over the number of memories, and so more memories down-weights attention probabilities. We want the attention vector to not be affected by the number of elements in the set. In particular, we weight the softmax by *log*(*n*_*memories*_).

#### Adding memories in WM models

There is no explicit memory adding process in WM models. There are simply input weights from the input at each timestep, ***i***_*t*_, and so the RNN must figure out how to hold these inputs in its recurrent memory.

#### Normalisation

We use normalisation in two places. 1) In EM models, after retrieving a memory i.e., before applying 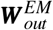 we perform a layer normalisation^60^: LayerNorm 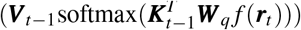. This is so all retrieved memories have the same scaling. 2) In some simulations (detailed in Table 2) we also normalise the output of the RNN for both WM and EM models. We do this for consistency when comparing WM and EM models. For WM models this means 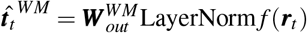. For EM models this means memories are retrieved as LayerNorm 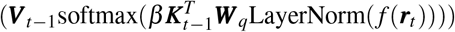.

**Table 2.** Hyperparameters for Figures in main text. Where there are commas, this indicates that multiple values were used for this hyperparameter, which will be explained in the individual figure. gRNN/rRNN/bRNN: GroupRNN/RegularRNN/BioRNN. X→Y means this hyperparameter was slowly changed from X to Y during training.

| Hyperparameter | Figure 2 | Figure 3 | Figure 4 | Figure 5 | Figure 6 |
| --- | --- | --- | --- | --- | --- |
| # RNN neurons | 32,64,128 | 256 | 512 | 128 | 128 |
| Activation Function | None | None,tanh,ReLU | ReLU | ReLU | ReLU |
| RNN type | gRNN | gRNN, rRNN, bRNN | rRNN | rRNN | rRNN |
| LayerNorm | True | False | True | False | False |
| Key/query dim | 64 | 64 | 64 | 64 | 64 |
| RNN regularisation | 0 | 0, 0.01 | 0 | 3e-4 | 3e-4 |
| KL regularisation | 0 | 0, 0.01 | 0 | 0 | 0 |
| Weight regularisation | 5e-8 | 1e-6 | 5e-8 | 1e-6 | 1e-6 |
| Batch Size | 128 | 128 | 128 | 128 | 128 |
| Optimizer | Amsgrad | Adam | Amsgrad | Adam | Adam |
| Learning Rate | 8e-4 → 2e-4 | 8e-4 → 2e-4 | 8e-4 → 2e-4 | 8e-4 → 2e-4 | 8e-4 → 2e-4 |
| Gradient steps | 100000 | 500000 | 100000 | 100000 | 100000 |
| # observations | 10 | 10 | 10 | 6 | 4 |
| Sample with replacement | True | True | False | False | False |

#### Optimisation

All weight matrices are learned by backpropagation in both models. We optimise a prediction loss on the target, *L*_*t*_, which is a cross-entropy loss when observations and targets are one-hot (all simulations other than Figures 5,6), or a squared error loss otherwise (Figures 5,6). For all simulations we add weights decay. For the ‘additional constraints’ in Figure 3 we add a L2 loss on the RNN activity. Also in Figure 3, we use an additional KL loss (where specified) which is a squared error loss between inferred (***h***) and predicted (*f* (***r***)) RNN activity. All loss hyperparameters specified in Table 2.

### A.3 Additional Task Details

Here we provide further task details. In all tasks we train on many instantiation of the task, i.e., observations randomised across each instantiation. In all tasks observations are randomly sampled at positions with replacement, except for the delay task and the PFC tasks.

#### A.3.1 Figures 2, 3, 11, 12, 13

##### Immediate Serial Recall (ISR)

Observations repeat cyclically, and the task is to predict the upcoming observation (Figure 1). There is no velocity signal. e.g., a 4-cycle: ***o*** = {1, 4, 3, 1, 4, 3, 1 …} and ***t*** = {−, −,−, 1, 4, 3, 1 …}. ‘−’ means a target that we do not try to predict since it has not been observed before and therefore is not possible to predict. The length of the each sequence is 7 multiplied by the repeat length.

##### N-Back

Observations are randomly drawn at each step, and the task is to recall the observation *n* timesteps ago. There is no velocity signal. e.g., for *N* = 3: ***o*** = {1, 2, 4, 2, 3, 5, 9 …} and ***t*** = {−,−, −, 1, 2, 4, 2…} The length of the each sequence is 7 multiplied by *N*.

##### 1D Navigation

Observations come from random walks on a loop. Velocity signals are provided. e.g., a 3-loop: ***o*** = {1, 3, 2, 2, 3, 1, 2 …},, ***v*** = {+1, +1, 0, 0, −1, −1, −1 …} and ***t*** ={−, −,−, 2, 3, 1, 2…} While velocities are randomly sampled at each step, to encourage exploration of the full loop, we increase the likelihood of repeating velocities (apart from the 0 velocity). The length of the each sequence is 7 multiplied by the loop size.

##### 2D Navigation

Observations come from random walks on a 2D torus. Velocity signals are provided. e.g., a 2x2 torus: ***o*** = {5, 1, 5, 2, 3, 1, 1 … }, ***v*** = {(+1, 0), (+1, 0), (0, +1), (−1, 0), (0, −1), (0, 0),… }, and ***t*** = {−, −, 5, −, −, 1, 1 …}. The length of the each sequence is 7 multiplied by the torus size.

#### A.3.2 Figure 4

##### 2-ISR with delays

Delay period lengths were randomly sampled from [3,4,5,6,7] for each task. Example observation sequences are ***o*** = {4, *, *, 2, *,*, 4, *, *, 2, …} or ***o*** = {1, *, 3, *, 1, *, 3, … } where * is the common delay observation (that also must be predicted). Corresponding target sequences are ***t*** ={−, −, −, −, −, −,−, *, *, 2, … } or ***t*** = {−, −,−, −, −, *, 3, Again we do not train on targets not possible to predict (i.e. observations/delays that have never been visited). We also so not train on the 1st return to the initial position. This is because it also cannot be predicted as it could have been a delay step. Observations are sampled without replacement. The length of the each sequence is 7 multiplied average full loop length (averaging over delays).

##### 4-ISR with delays

Delay period lengths were randomly sampled from [3,4,5] for each task. Example observation sequences are ***o*** = {4, *, 2, *, *, 2, *, 6, 4, *, 2, *, *, 2,, 6, … } or ***o*** = {1, 3, *, 8,, 2, 1, 3, *, 8, *, 2, … } where * is the common delay observation (that also must be predicted). Corresponding target sequences are ***t*** = {−, −, −,−, −, −, −,−, −, *, 2, *, *, 2, *, 6, … } or ***t*** ={−, −, −, −, −, −, −, 3, *, 8, *, 2, … } Again we do not train on targets not possible to predict (i.e. observations/delays that have never been visited). We also so not train on the 1st return to the initial position. This is because it also cannot be predicted as it could have been a delay step. Observations are sampled without replacement. The length of the each sequence is 7 multiplied average full loop length (averaging over delays).

#### A.3.3 Figure 14

##### 1D Navigation with structured velocity signals

We used a 7-loop task (1D navigation), but where the underlying velocity cycles a fixed schedule of {+2, +1, 0, − 1, 0, +1, +2, +1, 0, … } (velocity cycled on a 6-loop). Velocity signals were not provided. The length of the overall sequence is 7 multiplied by the loop size.

##### 2D Navigation with structured velocity signals

Sequences we drawn from a 3-by-3 torus, but where the underlying velocity cycles a fixed schedule of {*N,W, S, S, E, E, N, N, N,W, S, S*, … } (velocity cycled on a 8-loop). Velocity signals were not provided. The length of the overall sequence is 7 multiplied by the torus size.

##### 1D Navigation using macro-actions

We used a 8-loop task (1D navigation). We used 9 possible macro actions: {[+1, +1, +1, +1], [+1, 0, +1, +1], [+1, +1, +1, −1], [−1, +1, 0, +1], [+1, +1, −1, −1], [0, −1, +1, −1], [−1, 0, 0, −1], [−1, −1, −1, 0], [−1, − 1, −1, − 1] }. Macro-actions were sampled randomly, with a slight bias to repeating the same macro-action (to encourage full exploration of the loop). The length of the overall sequence is 7 multiplied by the loop size.

##### 2D Navigation using macro-actions

Sequences we drawn from a 3-by-3 torus. We used 8 possible macro actions: {[*N,W*, 0, *E*], [*S*, 0, *E, N*], [*N, S, E*, 0], [0, *S,W, E*], [*S, S, E, N*], [*W,W, S, E*], [*N, N,W, S*], [*E, E, N,W*]] }, where 0 means no velocity. Macro-actions were sampled randomly. The length of the overall sequence is 7 multiplied by the torus size.

#### A.3.4 Figure 5

**Xie et al**. Here observations are one of six 2D positions lying on a ring. This is just like the original paper^29^). Thus the observation vector consists of two real numbers rather than a one-hot vector as before. Likewise for the targets. Observations are sampled without replacement. Otherwise the set-up is the same as a 3-ISR task, but only with a single repeat (i.e. 6 observations in total), e.g., {***p***_5_, ***p***_3_, ***p***_2_, ***p***_5_, ***p***_3_, ***p***_2_ } and ***t*** = {−, −, −, ***p***_5_, ***p***_3_, ***p***_2_ }where ***p***_*i*_ is a 2D position vector (corresponding to the six positions lying on a ring).

#### A.3.5 Figure 6

**Panichello et al**. Here observations are one of four 2D coordinates lying on a ring. The original paper uses randomly sampled colours (from a 2D colour ring), and for visualisation purposes they bin these colours into 4 bins. Thus we just use 4 example colours (we could have used randomly drawn colours - the activity slot predictions are the same). Thus the observation vector consists of two real numbers (corresponding to a colour the colour wheel) rather than a one-hot vector as before. Likewise for the targets. Observations are sampled without replacement. The length of the sequence is 4 (top colour, bottom colour, cue, target), e.g., {***c***_3_, ***c***_2_, *cue* = *bottom*, *} and ***t*** = {−, −, −, ***c***_2_}, where ***c*** is a 2D vector (corresponding to a colour on the colour wheel).

### A.4 Analysis Details

#### A.4.1 Decoding Analyses

For all decoding analyses, we take the neural representations at each timestep in the sequence, along with the value of the variable we wish to decode at each timestep in the sequence. We collect this data for many timesteps and for multiple (240) random sequences, and collate this data into a training dataset (number of data-points is 240 *× s* where *s* is sequence length), with inputs being the neural representations, and outputs being the variable we wish to decode. In all decoding analyses, we only decode from timesteps after all positions have been visited at least once (so activity slots can be filled). We use (multi-class) logistic regression (using the sklearn python package) for all decoding analyses (apart from task-progress slot decoding - see later), as we discretise continuous random variables into discrete classes. We train the decoders and then test on a held out dataset of 60 random sequences (all presented results are test accuracy). We now describe how we obtain the particular variables we wish to decode.

##### Position decoding

We define the first element of each sequence to be at position 0, and then track position from then onwards (i.e., we track relative position to initial position). We give each position a unique identifier, so we can perform multi-class logistic regression.

##### Slot decoding

For every sequence, we calculate each slot’s stored observation according to our theory at all timesteps (Figure 2H; we call this the slot ‘slot-sequence’ for each slot). We decode each slot-sequence individually using multi-class logistic regression. When taking the average decoding performance over slots, we do not include slot 1, i.e., the input slot, as it will always decode to 1 (since the slot-sequence for slot 1 is just in input sequence which gets provided to the model). While for each task the observations in the slots are permutations of one-another (e.g., Figure 2H) and so it may be thought that if you can decode slot 1 (the input slot!) then you can decode any other slot, the permutation is not consistent over task instantiations. Thus, since we decode multiple sequences from multiple task instantiations at once, only RNNs that have actually learned a slot representation will have high slot-sequence decoding accuracy.

##### Past decoding

For every sequence, at each timestep we calculate which observation occurred *n* timesteps ago (Figure 2H; we call this ‘past-sequence’ for each time lag *n*). We do this for *n* = 0 to *n* = *n*_*s*_ − 1 where *n*_*s*_ is the number of slots. We run a separate decoder for each value of *n*. When taking the average decoding performance over past-sequences, do not include *n* = 0 as it will always decode to 1 (since the current observation is being provided to the model).

##### Progress decoding

To calculate progress, at each timestep, we calculate the number of delay steps taken since an observation and divide by the total number of delay steps (between neighbouring observations). We then discretise progress into 4 chunks, i.e. 0% → 25%, 25% → 50%, 50% → 75%, 75% → 100%.

##### Progress-velocity decoding

We calculate progress-velocity, for each timestep, we compute 1 divided by the total number of delay steps (between neighbouring observations). We then provide a unique identifier to its value. Thus decoding progress-velocity, is the same as decoding the number of delay steps between the current neighbouring observations.

##### Task-progress slot decoding

We first discrete the task into *T × N* task-progress slots where *T* is positions in task (i.e., 2 for the 2-ISR delay task) and *N* = 4 is the number of progress chunks after discretising progress. At each timestep we calculate which observation our theory says should be in each task-progress slot. Most slots will be empty as there are only *T* observations but *T × N* task-progress slots. Thus rather than having a separate decoder for each task-progress slot, we train a single 2-layer linear neural network (with sigmoid activation function on the output; 2 linear layers as it trains faster even though it is no more expressive) where the inputs are neural representation at each timestep and the outputs are whether an observation is in a particular slot. There output dimension is then *n*_*o*_ *× T× N* = 80 for the 2-ISR delay task, where *n*_*o*_ is the number of observations. When observation *a* is task-progress slot *b*, then the corresponding output neuron is 1 and 0 otherwise. Thus most entries will be 0 for the 2-ISR delay task: there will only be two 1 entries out of *n*_*o*_*TN* = 80, and so 0.975 is the baseline accuracy (which we normalise to in Figure 4).

#### A.4.2 Mutual Information Ratio

To calculate the mutual information ratio^43^ (MIR), we first compute the mutual information between all active neurons (defined as neurons with average activity greater than 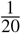 of the average neural activity) and the ‘slot-sequence’ for each slot (slot-sequences are the predicted stream of observations in each slot; Figure 2H). This gives a matrix of dimensions number of active neurons by number of slots. For each neuron we take its mutual information vector (of dimension number of slots), and calculate the maximum value of the vector divided by the sum of the vector (mutual information is always non-negative). This determines how specialised each neuron is for each slot. The MIR is then the average value over all active neurons. We do not include slot 1, i.e., the input slot, in this analysis.

#### A.4.3 Slot Algebra

To compute the slot algebra score, we generate sequences from four environments of identical structure, where the environments differ in their arrangement of observations in a special way. In particular the first environment has a random arrangement of observations, the second environment has a random arrangement of observations apart from one position (position *p*) where it has the same observation as the first environment. The third environment has the same arrangement of observations as the second environment apart from position *p* where it has a different, random, observation. The fourth environment is the same as the first environment, but in position *p* it has the same observation as the third environment. We then generate different random sequences for each of these environments. We choose a random position *i*, and record the neural representation for a random visit to position *i* on each of the four sequences. This gives us *Rep*_*i*_(*env*_1_), *Rep*_*i*_(*env*_2_), *Rep*_*i*_(*env*_3_), and *Rep*_*i*_(*env*_4_), where *Rep*_*i*_(*env*_*j*_) means the neural representation of environment *j* when at position *i*. The slot algebra score 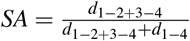, where (*d*_1−2+3−4_) is ||*Rep*_*i*_(*env*_1_) −*Rep*_*i*_(*env*_2_) + *Rep*_*i*_(*env*_3_) −*Rep*_*i*_(*env*_4_)||^2^ and *d*_1−4_ = ||*Rep*_*i*_(*env*_1_) −*Rep*_*i*_(*env*_3_)||^2^ where ||*…*|| ^2^ means square and sum all vector elements. We repeat this process 200 times for each model and average the individual *SA* scores to give the overall slot algebra score for that model.

We occasionally use an extra regularisation term (KL) which asks the ‘predicted’ internal representation, ***r***_*t*_, to the same as the ‘inferred’ representation, ***h***_*t*_. In particular, we add an extra term to the loss that is *λ*_*KL*_ ||***h***_*t*_− ***r***_*t*_|| ^2^, where *λ*_*KL*_ is the regularisation strength (see Table 2 for its value).

To ensure that the poor slot algebra scores of the regular RNN are not simply due to the fact that we provide the velocity of the *upcoming transition* to the network at time-step *t* (i.e. the recurrent input ***r***_*t*_ = ***W*** _*rec*_***h***_*t*−1_, and the external input, ***i***_*t*_, is ***o***_*t*_ and ***v***_*t*_ concatenated; reminder that ***v***_*t*_ is the velocity from that transition from ‘true underlying’ state ***z***_*t*_ to ***z***_*t*+1_), we additionally train networks where the velocity signal of the upcoming transition is only provided to the path integration step, i.e. ***r***_*t*_ = ***W*** _*rec*_***h***_*t*−1_ + *B****v***_*t*_ (***B*** is a learnable matrix), and the external input, ***i***_*t*_, is just ***o***_*t*_. This new network, in theory, could have no contamination of its internal representation with velocity, i.e., ***h***_*t*_ could be independent of velocity, and therefore have low slot algebra scores. Nevertheless, we find that these networks do not have low slot algebra scores (Figure 10), suggesting that the ***h***_*t*_ is dependent on the velocities signals taken to reach state ***z***_*t*_.

#### A.4.4 PFC Analyses

We followed the exact analyses described in Xie et al., 2022^29^ for Figure 5 and the exact analyses described in Panichello et al., 2021^33^ for Figure 6.

## Acknowledgements

We thank Kris Jensen and Jo Warren for helpful feedback on the manuscript. We thank the following funding sources: Sir Henry Wellcome Postdoctoral Fellowship (222817/Z/21/Z) to JCRW.; the Gatsby Charitable Foundation to WD.; Wellcome Principal Research Fellowship (219525/Z/19/Z), Wellcome Collaborator award (214314/Z/18/Z), and Jean-François and Marie-Laure de Clermont-Tonnerre Foundation award (JSMF220020372) to TEJB.; the Wellcome Centre for Integrative Neuroimaging is supported by core funding from the Wellcome Trust (203139/Z/16/Z); the James S. McDonnell, Simons Foundations, NTT Research, and an NSF CAREER Award to SG.

## Author Contributions

JCRW conceptualised the study in conversations with WD, TEJB, SG, ME. JCRW developed theory and performed simulations. JCRW wrote the manuscript with input from WD, TEJB, SG, ME.

## Materials & Correspondence

Correspondence to James CR Whittington.

## Data availability

No data was generated in this paper..

## Code availability

Python and Pytorch code will be made available on publication.

## Footnotes

i We note that our proposed WM activity slot solution may not the universal solution to all WM tasks. However we contend that for tasks involving non-trivial velocity signals, it is the solution neural networks learn, as evidenced by our subsequent simulations. See Discussion for further consideration.

ii It is important to consider each agent position *i* individually, since activity slot representations change from agent position to position.

iii We do not use the n-back dataset as there is no repetition of veridical position.

iv While these task-progress slots will now be continuous rather than discrete, with bumps of activity on continuous attractor manifolds now defining an observation within a ‘slot’, for consistency we still speak about them as if they were discrete.

v Normally key, query, and value matrices, are used i.e.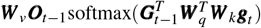. We leave them out for simplicity, but the following argument would remain the same if we included them.

vi If the target were to predict a mixture of past observations, the attention vector would not be one-hot, but the same argument follows just the same.

